# Short single-stranded DNA with putative non-canonical structures comprises a novel class of plasma cell-free DNA

**DOI:** 10.1101/2021.05.25.445702

**Authors:** Osamu Hisano, Takashi Ito, Fumihito Miura

## Abstract

Cell-free DNA (cfDNA) in human blood is currently investigated as a minimally invasive, highly informative biomarker. Here, we aimed to investigate the existence of the shorter cfDNA fragments in the blood. Using an improved cfDNA purification protocol and a 3′-end-labeling method, we found DNA fragments of approximately 50 nucleotides in human plasma, present at a molar concentration comparable to that of the nucleosome-sized fragments. These short fragments cannot be recovered by widely used cfDNA isolation methods, and are composed of single-stranded DNA (ssDNA), thus escaping detection in previous studies. We established a library-preparation protocol based on our unique ssDNA ligation technique and applied it to the isolated cfDNA. Deep sequencing of these libraries revealed that the short fragments are derived from hundreds of thousands of genomic sites in open chromatin regions and enriched with transcription factor-binding sites. Remarkably, antisense strands of putative G-quadruplex motifs occupy as much as one-third of peaks called with these short fragments. Hence, we propose a novel class of plasma cfDNA composed of short single-stranded fragments that potentially form non-canonical DNA structures.

## INTRODUCTION

Cell-free nucleic acids, detected in our bodily fluids, are attracting intense attention as a diagnostic material. In particular, cell-free DNA (cfDNA) in the blood is being focused on as a promising biomarker that can be measured with minimal invasion (i.e., liquid biopsy) (Wan et al. 2017; Bronkhorst et al. 2019). For example, detecting fetal cfDNA in the mother’s blood is a current practice that enables safe prenatal diagnosis (Lo et al. 2010). Meanwhile, cell-free tumor DNA (ctDNA) has been intensively studied as a potential biomarker for cancer diagnosis and follow-up after treatment (Cohen et al. 2017; Bronkhorst et al. 2019). Additionally, cfDNA in recipients after organ transplantation is used to monitor adverse side effects, particularly the rejection of transplanted grafts (Burnham et al. 2017).

The half-life of cfDNA in the bloodstream is generally short. A model experiment that traced the fate of radio-labeled DNA injected into mouse bloodstream revealed its rapid clearance through the kidneys (Tsumita and Iwanaga 1963). Meanwhile, approximately 70% of radio-labeled nucleosomes are removed by the liver (Gauthier et al. 1996). Moreover, fetal cfDNA rapidly declines in the mother’s blood after delivery, leading to an estimated half-life within one hour (Yu et al. 2013). The liver and kidneys play major roles in the clearance of cfDNA from the bloodstream (Chused et al. 1972).

The origin of cfDNA is considered to be apoptotic dead cells in various organs (Aucamp et al. 2018). As a result of cell death, nuclear DNA is fragmented by nucleases and released into the blood (Watanabe et al. 2019; Han et al. 2020). The size of cfDNA generally ranges from 130 bp to 180 bp (Jahr et al. 2001; Chan et al. 2004). Based on this size range, cfDNA is assumed to reflect the nucleosome structure of the source cells (Jahr et al. 2001; Chan et al. 2004). In healthy individuals, the primary source of blood cfDNA is hematopoietic cells (Snyder et al. 2016). In the blood of patients with cancer, organ recipients, and pregnant mothers, the level of cfDNAs originating from the tumors, transplanted grafts, and fetuses, respectively, are elevated (Lo et al. 2010; Burnham et al. 2017; Cohen et al. 2017; Bronkhorst et al. 2019). These cfDNAs can be distinguished from physiological DNA based on genetic variations, including mutations and single nucleotide polymorphisms. Recently, the epigenetic status of cfDNA has attracted intense attention. The epigenome differs from one cell type to another. Even in a single cell type, it varies depending on the cellular state. Accordingly, epigenetic information should extend the diagnostic capability of cfDNA to diseases that are not associated with genetic variations. DNA methylation is useful for this purpose because of its stability (Sun et al. 2015; Lehmann-Werman et al. 2016). Similarly, the fragmentation patterns of cfDNA may also provide valuable information as they reflect the chromatin status of the cells (Cristiano et al. 2019; Sun et al. 2019).

Next-generation sequencing (NGS) has substantially served to the advancement of cfDNA research. Specifically, deep sequencing by NGS enables the detection of genetic variations in a limited fraction of cfDNA, as well as changes in DNA methylation and fragmentation patterns. To read the nucleotide sequence of a DNA fragment using NGS, the fragment must be connected to two different adapters at either end. T4 DNA ligase-based protocols have also been widely applied for the analysis of cfDNA. Since T4 DNA ligase is active only on double-stranded DNA (dsDNA), it cannot be used for the adapter tagging of single-stranded DNA (ssDNA) unless a specialized adapter is introduced. Therefore, while there are plenty of publications investigating double-stranded cfDNA in the blood, studies focused on single-stranded cfDNA have remained limited until recently. However, with the advent of library preparation methods for ssDNA (Gansauge and Meyer 2013; Gansauge et al. 2017; Wu and Lambowitz 2017). the characterization of single-stranded cfDNA in the blood has begun to appear in the literature (Burnham et al. 2016; Snyder et al. 2016).

The existence of short cfDNA smaller than the nucleosome-sized cfDNA present in the blood has been previously described (Mouliere et al. 2011; Leszinski et al. 2014; Sanchez et al. 2018). Considering that apoptotic cells produce nucleases (Aucamp et al. 2018; Watanabe et al. 2019; Han et al. 2020) and that the blood presents nuclease activities (Watanabe et al. 2019; Han et al. 2020), it is likely that the cfDNA would get exposed to such nuclease activities and subsequently damaged, even if they were protected in the nucleosome structure. The extent of cfDNA damage has been related, at a certain extent, to the health condition of a person, and earlier studies have shown that the integrity of cfDNA becomes lower in patients with tumors (Mouliere et al. 2011; Leszinski et al. 2014; Sanchez et al. 2018). The ssDNA-adapted library preparation methods are effective to detect such damages in cfDNA as these methods can identify breakpoints inside the nucleosome-sized dsDNA (Burnham et al. 2016; Snyder et al. 2016; Sanchez et al. 2018).

Here, we investigated short cfDNA in human plasma using an improved cfDNA purification protocol and an advanced method for NGS library preparation from ssDNA. Consequently, we identified an abundant albeit heretofore overlooked class of cfDNAs that was notably composed of short ssDNAs enriched for characteristic sequences that potentially form non-canonical structures.

## RESULTS

### Identification of short ssDNAs in the cell-free fraction of blood

While there are plenty of reports analyzing cfDNAs of approximately 160 nucleotides (nt) in length, studies on shorter cfDNA fragments, especially those that combined advanced sequencing technologies, are limited. Therefore, we investigated whether, and to what extent, DNA fragments smaller than 120 nt exist in the cell-free fraction of human blood. We first used commercially available kits for cfDNA isolation from blood. However, these kits yielded poor recovery of short DNA fragments. As shown in Supplementary Figure S1A, all the kits failed to recover synthetic oligodeoxyribonucleotides (ODNs) shorter than 60 nt, even when using the optional protocols for short nucleic acids provided by the manufacturers. We thus used a traditional or conventional DNA isolation method that employs <u>p</u>roteinase K treatment, <u>p</u>henol-chloroform extraction, and <u>i</u>sopropanol <u>p</u>recipitation (PPIP method). As shown in Figure 1A, the PPIP method enabled quantitative recovery of ODNs as short as 30 nt.

**Figure 1.**
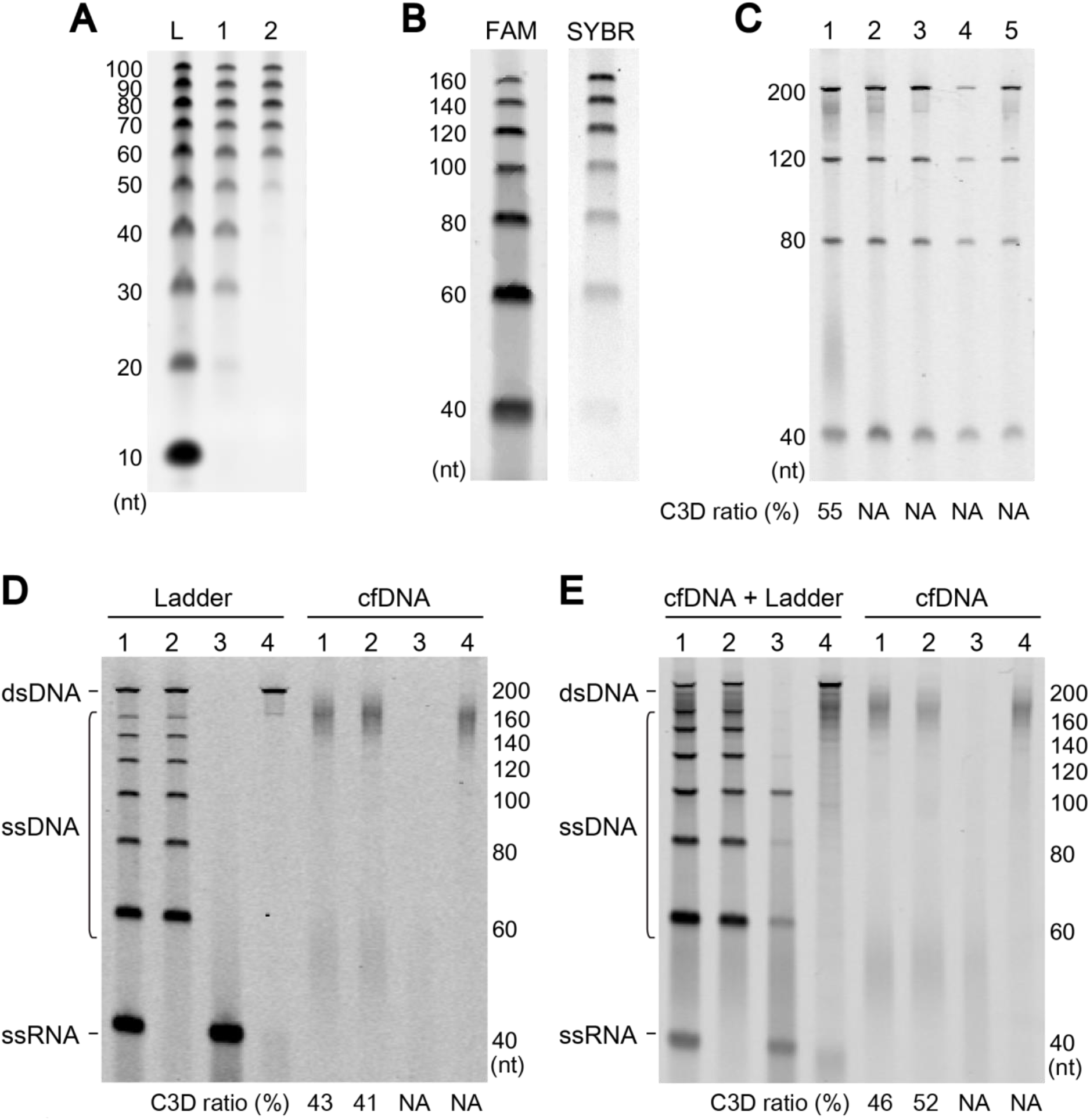
Short single-stranded DNA in the cell-free fraction of blood. **A.** Recovery of 10 bp DNA step ladder (lane L, Promega, Fitchburg, WI, USA) was compared using the PPIP method (lane 1) and QIAamp Circulating Nucleic Acid Kit (lane 2). nt; nucleotides **B.** An ODN mixture was detected with 3′-terminal fluorescent labeling (FAM) or staining with SYBR Gold nucleic acid stain (SYBR). C. cfDNA purified from the same plasma using the PPIP scheme or four commercially available kits were compared. The purification methods are the PPIP (lane 1), the QIAamp Circulating Nucleic Acid Kit from Qiagen (Hilden, Germany) (lane 2), the Plasma/Serum Cell-Free Circulating DNA Purification Mini Kit from Norgen Biotek (Thorold, Canada) (lane 3), the NucleoSpin Plasma XS from Takara Bio Inc. (Shiga, Japan) (lane 4), and the NEXTprep-Mag cfDNA Isolation Kit from PerkinElmer (Waltham, MA, USA) (lane 5). **D, E.** Effects of nuclease treatments on a model nucleic acid mixture and cfDNA. The treatment was performed for purified cfDNA (D) or in the plasma before purification (E). The nucleases used were as follows: no enzyme (lane 1); E. coli ribonuclease I (lane 2); DNase I (lane 3); *E. coli* exonuclease I and Rec J (lane 4). For the internal control in E, a model DNA mixture was spiked into the plasma (left). For details, see Supplementary Methods.

The signal intensity of DNA stained with intercalating dyes, such as ethidium bromide and SYBR Gold, is dependent on DNA mass. Accordingly, signal intensity per DNA molecule reduces proportionally to the size of DNA. Hence, the low signal intensity of SYBR Gold-stained short ODNs suggested a potential risk for short cfDNAs escaping detection using conventional gel staining. To circumvent this risk, we used terminal deoxynucleotidyl transferase (TdT) to label the 3′-end of each DNA molecule with a fluorophore-bearing nucleotide and detected the fluorescence after gel electrophoretic separation. Since the signal detected with this procedure accurately reflects the copy number, not the size of the DNA molecule, the signal-to-noise ratio should be improved, particularly in the small DNA range. This strategy drastically improved the sensitivity for detecting short ODNs compared to the intercalating dye-based method (Figure 1B).

We analyzed the DNA extracted from the cell-free fraction of human blood by combining the PPIP method with the 3′-end labeling. The results revealed that DNA fragments of ~50 nt were abundantly present in both plasma and serum of all the donors (Figure 1C, lane 1; and Supplementary Figures S1B and C). On the other hand, we could not detect any signal around 50 nt for the cfDNA purified with any of the commercially available kits (Figure 1C, lanes 2-5, Supplementary Figure S1D). Importantly, the molar amount of these short DNA fragments is comparable to nucleosome-sized ones (Figure 1C, lane 1, Supplementary Figure S1D). Both the short and nucleosome-sized fragments were resistant to *E. coli* ribonuclease I; therefore, they should be DNA (Figure 1D and E, Supplementary Figure S1E and F). Interestingly, the short DNA fragments were sensitive to ssDNA-specific exonuclease treatment, whereas the nucleosome-sized fragments were not (Figure 1D and E, Supplementary Figure S1E and F), suggesting that most of the short fragments are composed of ssDNA. In the present study, we call the short DNA fragment C3D as an abbreviation of <u>c</u>ell-free short single-stranded (<u>3</u>S) <u>D</u>NA and the ~160-nt DNA fragment as nucleosome-protected DNA (NPD).

### C3D exists in the liquid phase, not in membranous vesicles in plasma

Since C3D is a short ssDNA and ssDNA is generally more labile than dsDNA, we sought to determine why C3D is abundantly found in plasma that has nuclease activities. It is well known that exosomes (small vesicles) contain nucleic acids that are protected from nuclease activities in the blood (Pos et al. 2018). We thus investigated whether C3D is found in the exosomes collected by ultracentrifugation of plasma and serum. While we were able to successfully enrich the exosomal fraction by ultracentrifugation at 100,000 ×*g* to detect the exosome-specific marker CD9 (Supplementary Figure S2A), we could not find any C3D signal in the exosome-enriched fraction (Supplementary Figure S2B). In addition, we detected DNA in the exosomal fraction (Supplementary Figure S2C), which is in accordance with previous reports (Kahlert et al. 2014; Thakur et al. 2014), but its size was quite different from that of C3D. These results strongly suggested that C3D exists in the liquid phase and not in the membranous vesicles. To verify these results, we treated the plasma with *E. coli* exonuclease I and found that the exonuclease treatment of plasma led to the disappearance of C3D, as did the treatment of purified cfDNA (Figure 1E). Thus, we concluded that C3D exists as a naked form or in a state susceptible to exonuclease digestion, in the liquid phase of blood.

### Strategy for C3D sequencing using highly efficient ssDNA ligation

Recently, we developed a highly efficient technique, termed TACS ligation, for adapter tagging of ssDNA (Miura et al. 2019). This technique comprises two successive enzymatic reactions. The first is TdT-mediated modification of the 3′-end of target ssDNA with a few adenylates (rAMP), and the second is RNA ligase-mediated adapter ligation to the modified 3′-end of the target DNA (Figure 2A and Supplementary Figure S3A). Notably, TACS ligation can ligate a 5′-phosphorylated adapter to the 3′-end of target ssDNA with more than 80% efficiency (Miura et al. 2019). We thus designed a scheme for library preparation from ssDNA based on TACS ligation (TACS-T4 scheme; Figure 2A). Since the product of TACS ligation contains a few rAMPs between the target DNA and the adapter, reverse transcriptase activity is required to synthesize DNA complementary to the adaptor-tagged ssDNA. We previously found that Taq DNA polymerase, and its mutants, exhibit such an activity that efficiently converts ssDNA with short RNA stretches to dsDNA (Miura et al. 2019).

**Figure 2.**
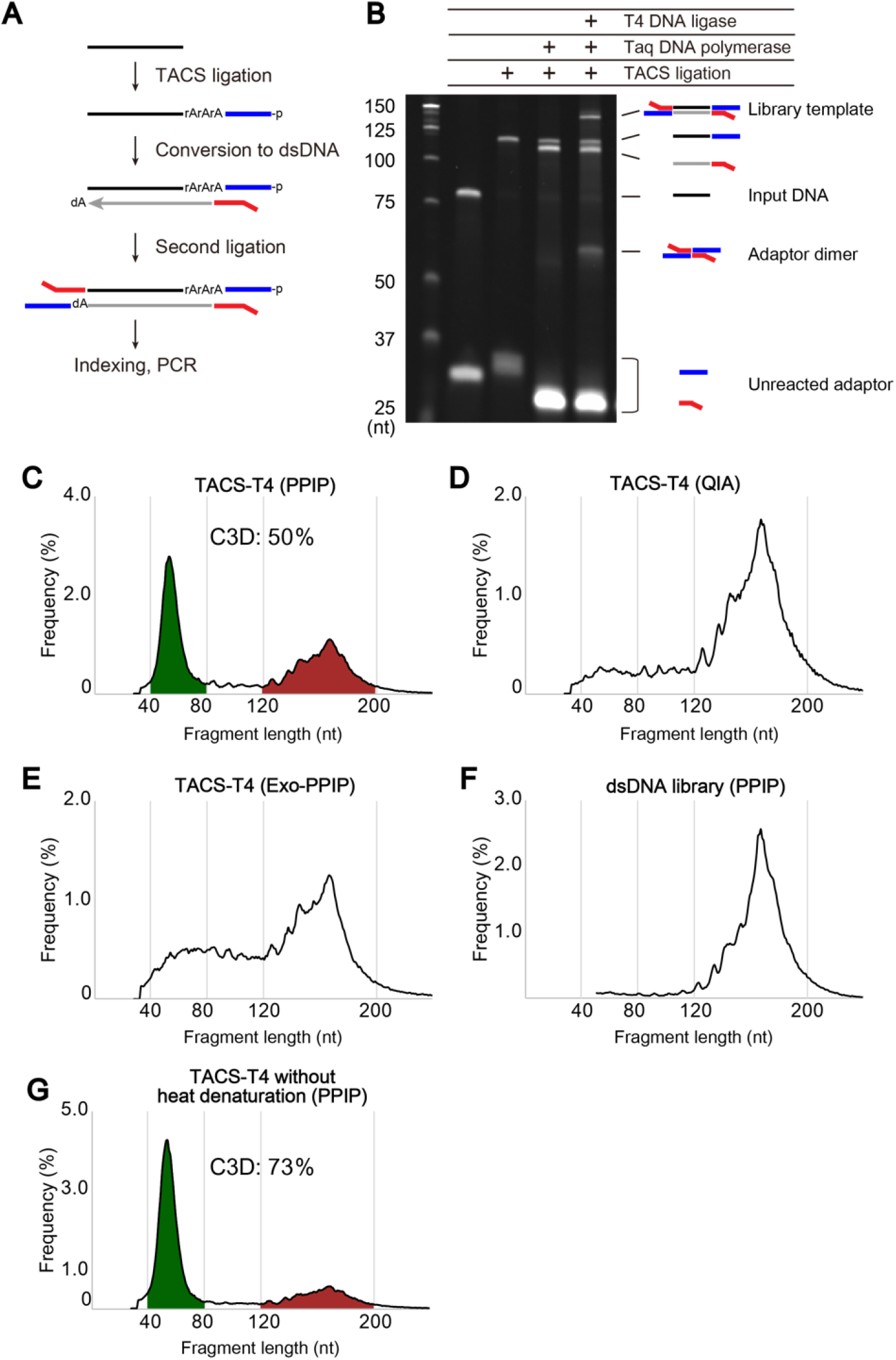
Library preparation from cfDNA. **A.** TACS-T4 scheme for library preparation from ssDNA. For details, see Supplementary Figure S3A. **B.** Efficiency of each step of TACS-T4 scheme applied to a model ODN. **C–G.** The size distribution of reads for five different cfDNA libraries. The cfDNAs used for the library preparations were purified with the PPIP method (C, E, F, and G) and QIAamp Circulating Nucleic Acid Kit (D). The PPIP purified DNA was treated with (E) or without (C, D, F, and G) a mixture of *E. coli* exonuclease I and Rec J before the library preparation. The heat denaturation before the library preparation was omitted for G. The libraries were prepared with TACS-T4 (C, D, E, and G) or a commercially available kit (F, ThruPLEX DNA-Seq kit, Takara Bio Inc.). For details, see Supplementary Methods.

Once fully double-stranded, T4 DNA ligase can attach a dsDNA adapter to the opposite end of the DNA (Figure 2A and Supplementary Figure S3A). In the TACS-T4 scheme, the dsDNA adapter is formed between the ssDNA adapter used for TACS ligation and the primer used for complementary DNA synthesis, after which it is ligated to the opposite end of the target DNA (Supplementary Figure S3A). Thus, the TACS-T4 scheme recycles the ODNs used in the previous steps to obviate the need for a purification step for their removal, thus aiming to improve the library yields.

Note that the TACS-T4 scheme includes an aprataxin-mediated deadenylation step to ensure the recycling strategy. RNA ligases first adenylate the phosphate group at the 5′-end of the donor and then connect the activated phosphate group to the 3′-hydroxyl end of the acceptor (Supplementary Figure S3A). Here, we observed that the 5′-phosphate group of the adapter was almost completely adenylated during the TACS ligation step and that this 5′-adenylation inhibited the second ligation step by T4 DNA ligase (Supplementary Figures S3B and C). Therefore, to remove the adenylate from the 5′-end of the adapter, we introduced an aprataxin-mediated deadenylation step, which enhanced the efficiency of the second ligation step (Supplementary Figures S3B and C).

Following the second adapter ligation and subsequent purification, the TACS-T4 scheme uses polymerase chain reaction (PCR) with DNA polymerase lacking reverse transcriptase activity to amplify and index the library. Note that the DNA polymerase can use only the DNA strand complementary to the input ssDNA. Therefore, the reads obtained by this method have strand specificity reflecting the ssDNA insert (Supplementary Figure S3D). Based on the results for the model experiment using a synthetic ODN, the implemented TACS-T4 scheme appeared to be efficient (Figure 2B) with approximately 20% of ODN converted to the library molecule (Supplementary Figure S3E).

### C3D is abundant and derived mainly from nuclear DNA

Next, we used the TACS-T4 scheme to prepare sequencing libraries from cfDNA isolated using the PPIP method. Since the TACS-T4 scheme preferentially converts ssDNA fragments to library molecules, cfDNA was heat-denatured prior to library preparation. We conducted a small-scale, paired-end sequencing of the library on Illumina MiSeq and calculated the end-to-end distance of the mapped paired-end reads on the reference genome. The size distribution of the library fragments formed two major peaks (Figure 2C, Supplementary Figures S4A, and S4B), one at approximately 160 nt and the other around 50 nt, consistent with the gel electrophoresis results for 3′-labeled cfDNA (Figures 1C–E and Supplementary Figures S1D–F). The former and the later fragments undoubtedly correspond to NPD and C3D, respectively.

In contrast, when a commercially available kit was used for cfDNA isolation, the peak appearing at 50 nt disappeared (Figure 2D and Supplementary Figure S4C), which was caused by the ineffectiveness of most commercially available kits to recover such short DNA fragments (Figure 1C). Similarly, when exonuclease I treatment was performed before heat denaturation of cfDNA, the peak appearing near 50 nt (i.e., C3D), but not that near 160 nt (i.e., NPD), disappeared (Figure 2E and Supplementary Figure S4D). The T4 DNA ligase-based commercial library preparation protocol optimized for dsDNA also failed to obtain a peak near 50 nt, even when the PPIP method was used for cfDNA purification (Figure 2F and Supplementary Figure S4E). Conversely, when the step involving heat denaturation before TACS ligation was omitted, the peak corresponding to NPD was diminished, leading to a concomitant fractional increase in the C3D peaks (Figure 2G and Supplementary Figure S4F). These results were expected, as the majority of NPD are double-stranded (Figure 1D and 1E, Supplementary Figure S1E and F) and, hence, not amenable to TACS ligation unless denatured.

Since the TACS-T4 scheme is a novel method, it is possible that the C3D peaks were an artifact specific to this scheme. To examine this possibility, we compared TACS-T4 with two different techniques previously reported (Gansauge and Meyer 2013; Gansauge et al. 2017) using the same cfDNA prepared with the PPIP method (Figure 1C, lane 1). As shown in Supplementary Table S1, TACS-T4 outperformed the other two in terms of library yields. The distribution size of the amplified libraries and sequenced reads of the three methods were almost identical (Supplementary Figure S5). In addition, compared to the input DNA, C3D appeared to be slightly underrepresented in the TACS-T4 library but rather overrepresented in the other two libraries. This was presumably due to the effects of extensive removal steps for elimination of the adaptor dimers in the TACS-T4 protocol, which likely caused the loss of shorter fragments, and to the effects of high PCR cycles in the other two protocols, which could lead to an amplification bias against longer fragments (Supplementary Figure S5B). Despite these differences, all three libraries formed two major peaks of reads corresponding to the peaks of input DNA fragments revealed by gel electrophoresis (Supplementary Figure S5). Therefore, the existence of C3D was supported not only by the TACS-T4 scheme but also by the other library preparation methods.

Recently, several studies have reported methods for library preparation from ssDNA, some of which were applied to cfDNA. For instance, Burnham et al. described the existence of ssDNA in the plasma (Burnham et al. 2016). They prepared two cfDNA libraries, one with an ssDNA-adapted protocol (ssDNA-lib) and the other with a conventional protocol adapted only to dsDNA (dsDNA-lib). They found that mitochondrial and microbial sequences are enriched in ssDNA-lib as short fragments ranging from 40 to 60 nucleotides. Since the size of C3D is similar to that of mitochondrial and microbial DNA fragments described by Burnham et al., we next investigated whether C3D originates from mitochondria or microbes. First, the reads obtained with TACS-T4 in the current study and ssDNA-lib by Burnham et al. (Burnham et al. 2016) were mapped to human nuclear and mitochondrial genome sequences using the same analytical pipeline. Next, the unmapped reads were examined to determine whether they were mapped to bacterial genomic sequences using the same procedure as that used by Burnham et al. The size distribution of mitochondrial and microbial fragments in both libraries peaked around 60 nt and 40 nt, respectively (Figures 3A and C). Therefore, both studies observed fragments of similar sizes originating from the mitochondrial and microbial genomes. As shown in Figures 3B and D, 91% and 90% of the reads were mapped to the human nuclear DNA in the TACS-T4 library and ssDNA-lib, respectively; the fraction of reads mapped to the mitochondrial genome was 0.11% and 0.01% in TACS-T4 and ssDNA-lib, respectively. Similarly, the fraction of reads mapped to the microbial genomes was marginal in both libraries. Therefore, the majority of the C3D originated from the nuclear genome, not mitochondrial or microbial genomes.

**Figure 3.**
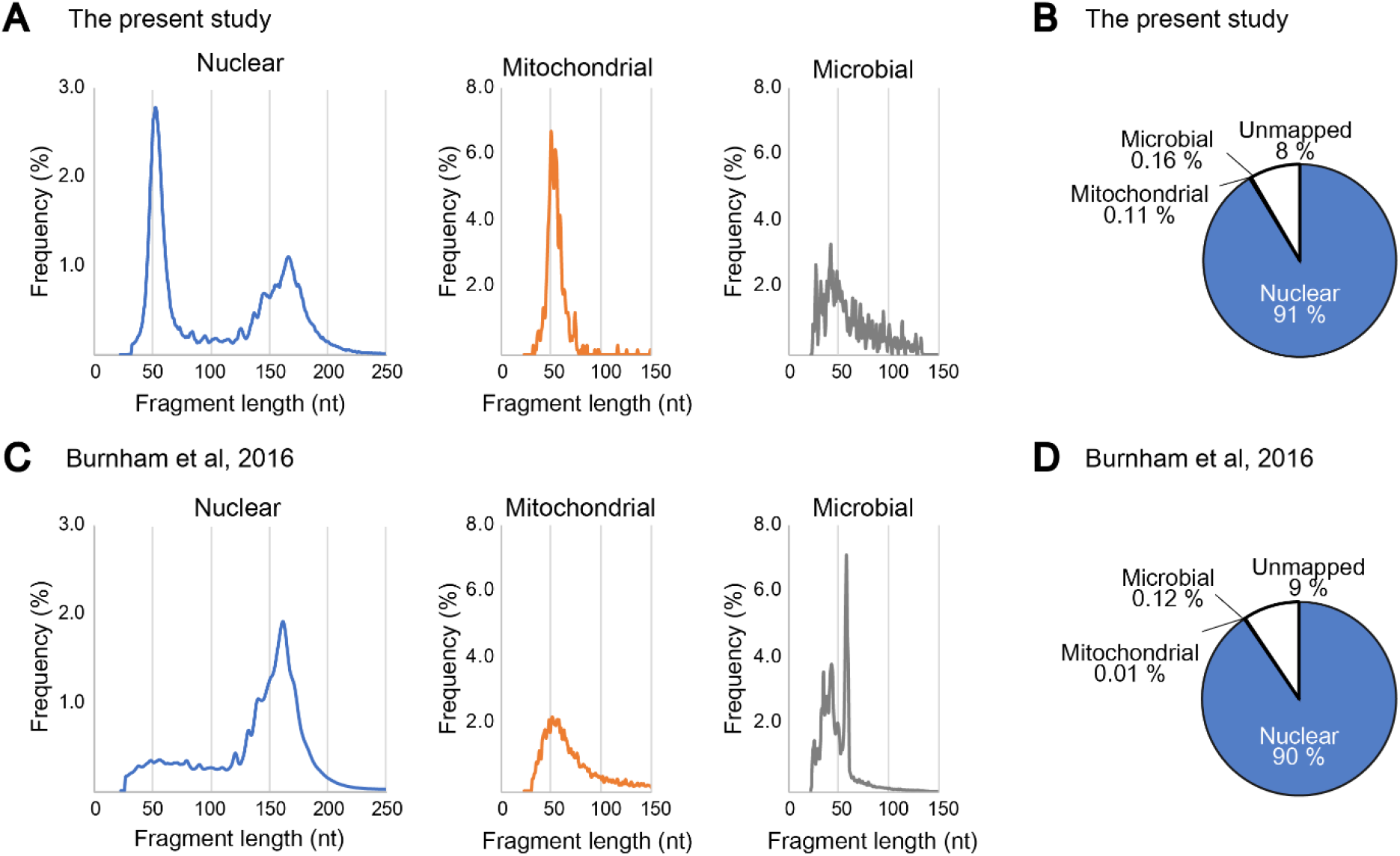
Most C3D is derived from nuclear DNA. **A.** The size distributions of cfDNA fragments for the library prepared with the TACS-T4 scheme using the cfDNA purified with the PPIP method are shown. Reads derived from nuclear (same data as in Figure 2C), mitochondrial, and microbial genomic DNAs were separately analyzed. **B.** Pie chart for the composition cfDNA reads in A. **C.** Size distribution of cfDNA fragments in Burnham et al. (2016). **D.** Pie chart for the composition cfDNA fragment in C. For details, see Supplementary Methods.

### Genomic origins of C3D are shared among individuals

We next prepared cfDNAs from five healthy individuals with the PPIP method, labeled their 3′-ends with a fluorophore, and separated them using denaturing polyacrylamide gel electrophoresis. As shown in Supplementary Figures S1B, S1C, and S6A, nearly the same patterns were shared by the five individuals, suggesting that C3D is generally present in healthy human blood. We then prepared sequencing libraries from these cfDNAs using the TACS-T4 scheme without heat denaturation to maximize the fraction of C3D reads. We sequenced the five libraries using HiSeq X, assigning one lane to each library (523M to 544M reads per sample, Supplementary Table S1), mapped the reads to the reference genome, and compared the distribution of C3D peaks among the five individuals. Manual inspection of the genome browser shots suggested that C3D peaks are distributed throughout the genome, and their positions were largely shared among the five individuals (Figure 4A). About one-fifth of C3D reads contributed to form these peaks (Supplementary Table S1). The total number of MAC2-called C3D peaks was largely comparable with approximately a hundred thousand (Figure 4B, Supplementary Table S1), and most of them were shared by at least two of the five individuals (Figure 4B and Supplementary Figure S6B). We also confirmed that the normalized read depths of individual peaks demonstrated good correlation among the five individuals (r = 0.91–0.95, Figure 4C; p-values = 0.13–0.49, Wilcoxon rank-sum test, Figure 4D). Taken together, C3D is likely generated from specific genomic loci with similar efficiency in any healthy individual.

**Figure 4.**
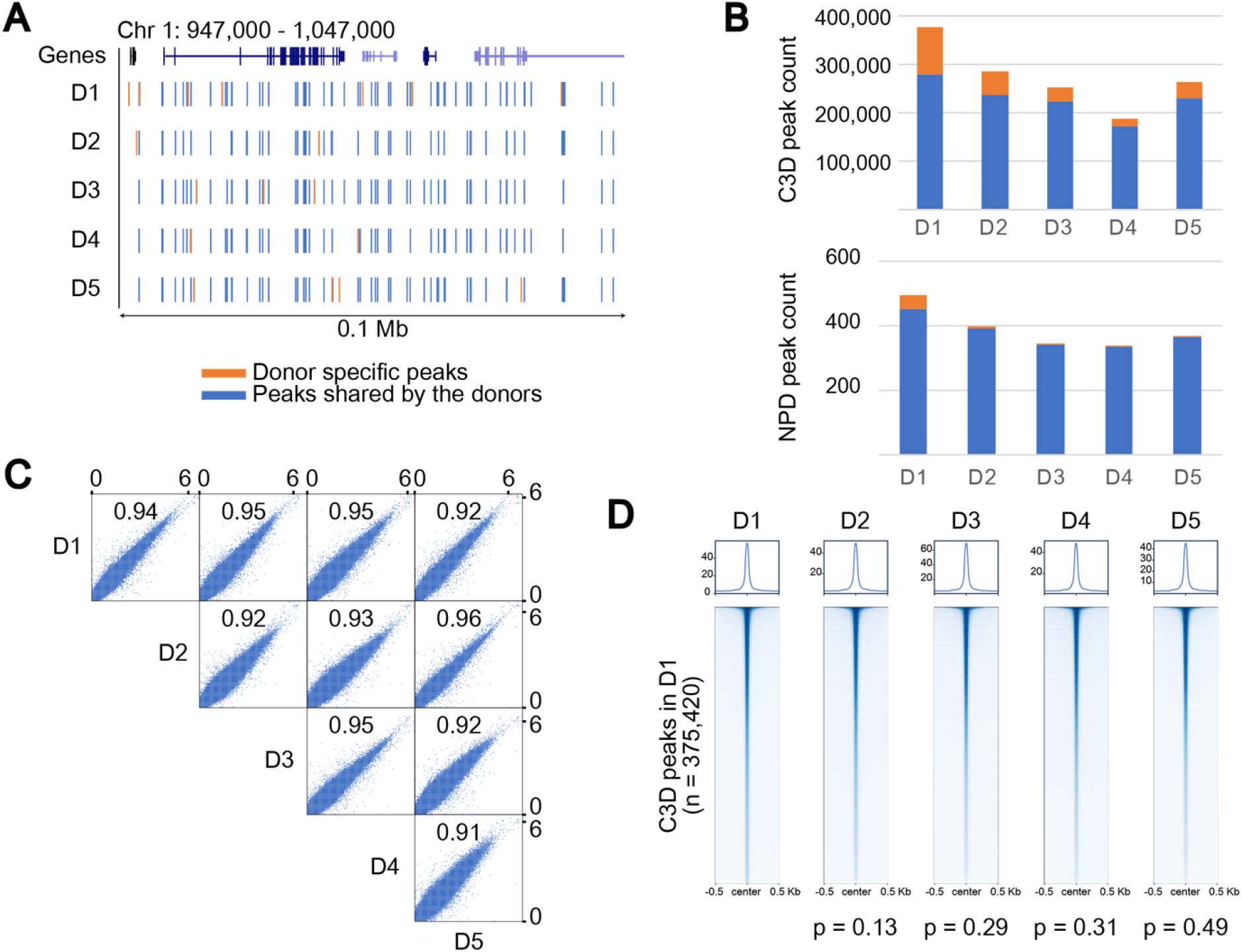
C3D peaks are shared among individuals. **A.** A browser shot comparing C3D peaks among five healthy donors. The peaks shared between at least two donors are shown in blue, whereas those specific for a certain individual are shown in orange. **B.** Total number of C3D and NPD peaks for each donor. Blue and orange indicate peaks shared by two or more donors and donor-specific peaks, respectively. **C.** A scatter plot matrix comparing the normalized read coverage of C3D peaks between two donors. The correlation coefficients are indicated in each plot. Both axes are on a base-10 logarithmic scale. **D.** Aggregation plots of the normalized read coverage for C3D peaks. C3D peaks are sorted according to the normalized read coverage in donor 1 (D1). This order is maintained in the aggregation plots for D2 to D5. The correlation coefficients obtained using the Wilcoxon rank-sum test are shown at the bottom.

### C3D is enriched in the regulatory regions of genes

To reveal the characteristics of C3D, we investigated the distribution of C3D peaks in the annotated genomic features. For this purpose, we used annotatePeaks of HOMER with the “Basic Annotation” provided in the package. The C3D peaks were found to be derived from all genomic features (Figure 5A). Interestingly, the C3D peaks appeared in 5′ UTRs and promoters at 3.6- and 3.5-fold higher than expected frequency, respectively (Figure 5B). Accordingly, an aggregation plot relative to the protein-coding genes formed a C3D peak in the promoter region (Figure 5C). Of the 59,461 protein-coding genes in RefSeqGene, 19,136 (34%) harbored C3D peaks in their promoter regions (Supplementary Figure S7A). Conversely, 2.9% of C3D peaks overlapped with promoters of protein-coding genes. Moreover, gene ontology analysis of genes with C3D peaks in their promoters and 5′ UTRs suggested that C3D is related to diverse functions (Supplementary Figure S7B). In contrast to the protein-coding genes, the C3D peaks were less enriched in the promoters of non-coding RNA genes (Supplementary Figure S7C). In addition to the “Basic Annotation,” the HOMER package provides “Detailed Annotation.” While no remarkable enrichments were observed in most of these annotations (Supplementary Figure S7D), strong enrichments were detected in regions annotated as CpG islands, low-complexity regions, and simple repeats (Figure 5D and Supplementary Figure S7D). Interestingly, an aggregation plot of C3D reads indicated that they were enriched at the boundary and flanking regions of CpG islands (i.e., CpG island shore) rather than within the CpG islands *per se* (Figure 5D). We performed the same enrichment analysis on the enhancers annotated by the FANTOM5 project (Andersson et al. 2014; Lizio et al. 2019) and found that the frequency of appearance of C3D peaks was 2.8 times higher than expected in these enhancers (Supplementary Figure S7D). It has been reported that the majority of the cfDNA in the blood of healthy individuals might have a hematopoietic origin (Snyder et al. 2016). Therefore, we investigated whether the C3D peaks are enriched in blood-specific promoters and enhancers. However, contrary to our expectations, the enrichment of C3D peaks was less prominent in the promoters and enhancers of blood-specific genes (Supplementary Figure S7E and F).

**Figure 5.**
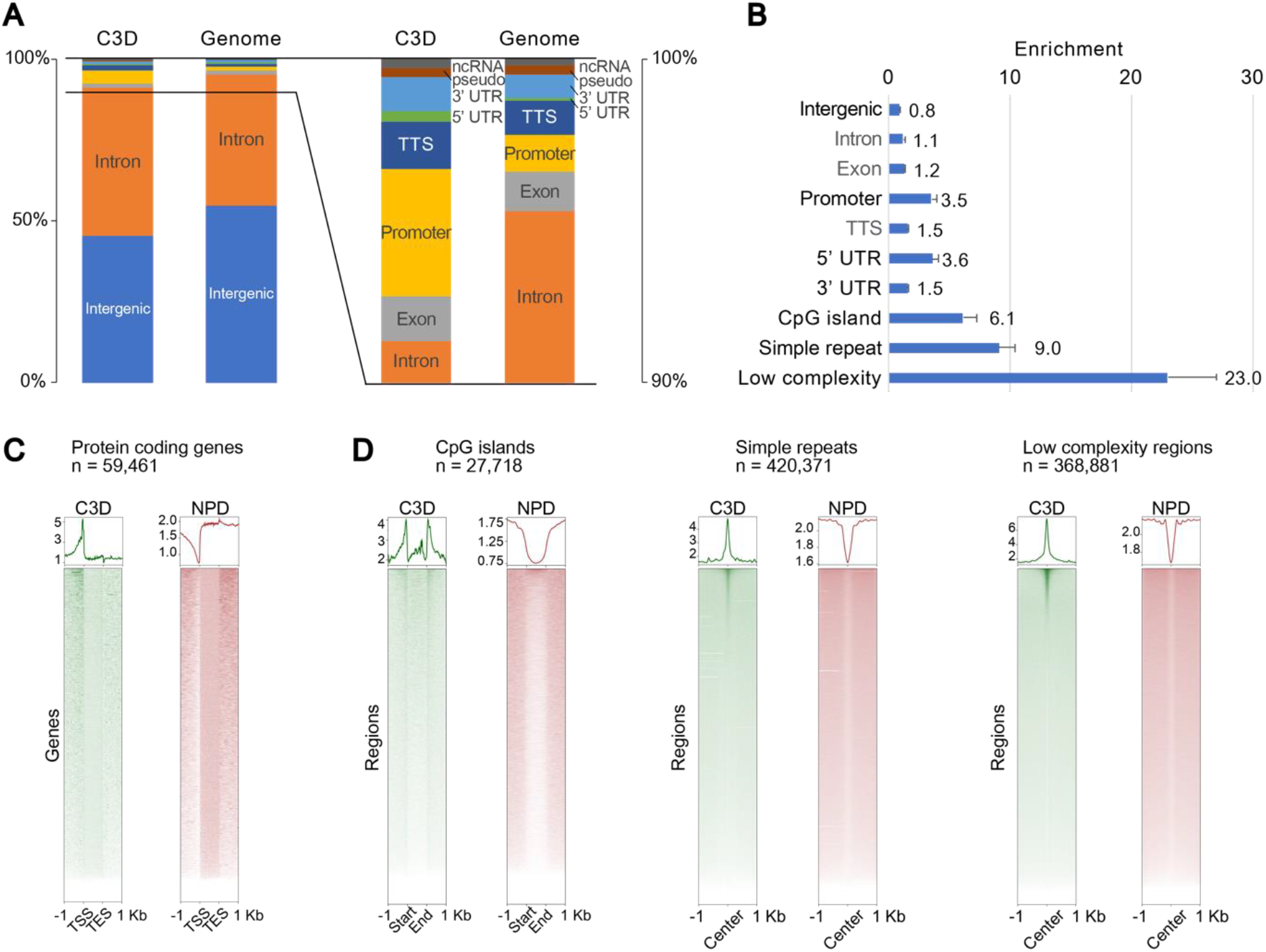
C3D is enriched in the regulatory regions of genes. **A.** Proportions of annotated genomic features assigned to C3D peaks (left) and the entire genome (right). **B.** Enrichment of annotations to C3D peaks over expected frequencies. For A and B, the average values of the five donors are shown. **C.** Aggregation plots of the normalized read coverage of C3D and NPD for protein-coding genes. **D.** Aggregation plots of the normalized read coverage of C3D and NPD for CpG islands, simple repeats and low-complexity regions.

### C3D colocalizes with the diverse functional features

The DNase I-hypersensitive sites (DHSs) and peaks of ATAC-seq colocalize with the promoter and regulatory regions of actively transcribed genes (Buenrostro et al. 2013). Knowing that C3D peaks are enriched in the promoter regions of genes, we next sought to determine whether DHSs and ATAC-seq peaks colocalize with C3D. As expected, C3D peaks were colocalized with both DHSs and ATAC-seq peaks (Figure 6). Although the colocalizations were statistically significant (p < 0.001, permutation test), only 3.8% and 5.3% of C3D peaks overlapped with DHSs and ATAC-seq peaks, respectively (Figure 6C), indicating that the origin of C3D peaks could not be explained solely by “open chromatin.” Recently, Snyder et al. (Snyder et al. 2016) and Burnham et al. (Burnham et al. 2016) revealed the presence of cfDNA shorter than 80 nt, which were specifically found in libraries prepared using a method adapted for ssDNA (Supplementary Table S2). These short cfDNAs colocalized with DHSs and binding sites of several transcription factors (TFs), including CCCTC-binding factor (CTCF) (Supplementary Figure S8). Thus, we extended the colocalization analysis to the ENCODE clustered TFs ChIP-seq data (Gerstein et al. 2012). We found that C3D peaks significantly colocalized with the binding sites of CTCF (Figure 6C) and other TFs; however, only a limited fraction of C3D peaks overlapped with the binding sites of individual TFs (TFBSs) (Supplementary Figure S9A–C, Supplementary Table S3). We further extended the comparison using the dataset downloaded from ChIP-Atlas (Oki et al. 2018) and confirmed that the overlapping of C3D peaks with the peaks annotated by other functional genomic analyses was statistically significant. However, the overlap was generally not very prominent (Supplementary Figure S9D–F, Supplementary Table S4).

**Figure 6.**
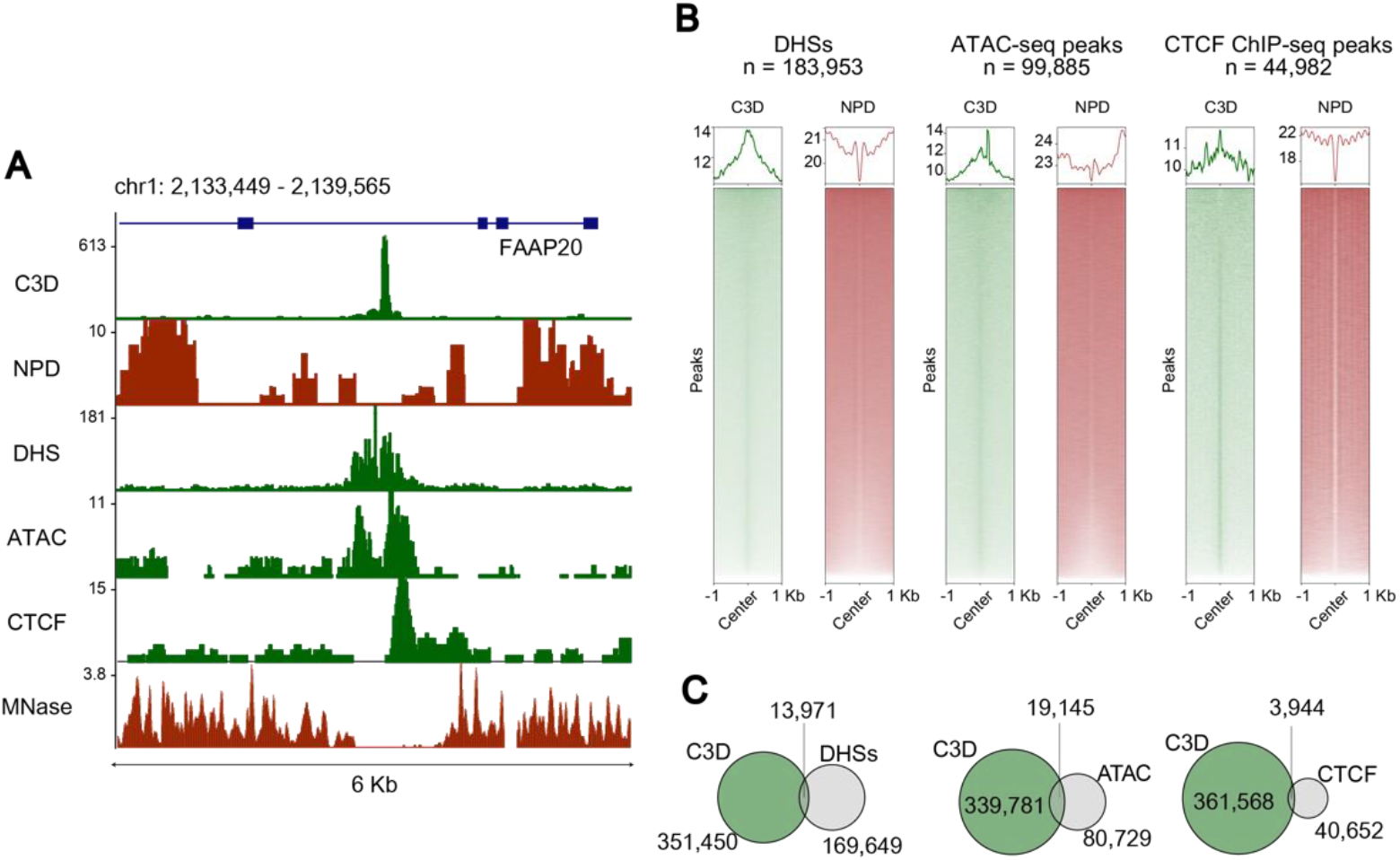
C3D colocalizes with DNaseI hypersensitive sites and several transcription factor-binding sites. **A.** A genome browser shot including tracks for C3D, NPD, DNase-seq (DNaseI hypersensitive sites, DHS), ATAC-seq, ChIP-seq for CTCF, and MNase-Seq. Data sources from the present study (C3D and NPD), ATAC-seq of GM12878 cells (Buenrostro et al. 2013), and ENCODE project (DHSs, CTCF, and MNase) (Rosenbloom et al. 2013). **B.** Aggregation plots for the read coverage of DNase-seq, ATAC-seq, and CTCF ChIP-seq for the C3D and NPD peaks. **C.** Venn diagrams showing the overlap of C3D peaks with DHS, the peaks of ATAC-Seq, and CTCF binding sites.

To know whether this small overlapping of the short cfDNA peaks with genomic features was a common characteristic, we performed peak calling with the data reported by Snyder et al. (Snyder et al. 2016) and Burnham et al. (Burnham et al. 2016). Surprisingly, the numbers of peaks called with these datasets (1,008 and 963 peaks for Snyder et al. (2016) and Burnham et al. (2016), respectively) were two orders of magnitude smaller than those of C3D (271,628 peaks) even after normalizing by the total number of reads (Supplementary Table S2). Despite the different number of peaks, similar trends were observed in the peaks called with the short cfDNA fragments; while we could recognize the enrichment of the peaks on TFBSs, the fractions of the peaks overlapping with the TFBSs were limited to less than one percent (Supplementary Figure S10 and Supplementary Table S5). Therefore, the majority of TFBSs are unrelated to the peaks of short cfDNA fragments.

The short cfDNA fragments presented good colocalization with TFBSs, whereas only a limited fraction of the cfDNA peaks overlap with TFBSs. These observations might be partially explained by the different localization trends of the peak-forming and the other reads. Only one-fifth of the C3D reads contributed to form peaks (Supplementary Table S1), which means that the remaining four-fifths did not contribute to peak formation. Then, we divided the C3D reads into two groups, those located on the peaks (C3D^on^) and those out of the peaks (C3D^off^), and separately analyzed them in aggregation plots. As expected, the patterns of the aggregation plots were largely different between the groups. We observed an enrichment of the C3D^off^ reads on the TFBSs, which is in accordance with that observed by Snyder et al. (2016) (Snyder et al. 2016), whereas the C3D^on^ reads were notably excluded from the centers of the TFBSs (Supplementary Figure S11). These results collectively demonstrated that C3D is composed by two groups: one is similar to previously described short cfDNAs, whereas the other is different and appears to be novel. Therefore, we have focused on the latter.

### Complementary strand of G4 motifs comprise one-third of C3D peaks

When inspecting the reads in the C3D peaks (i.e., C3D^on^), we found that many were extremely C-rich (Figures 7A and B), and contained simple repeats and low-complexity sequences, which were already exemplified as the enrichment of such genomic features (Figure 5B). Interestingly, the coverage of these C3D peaks exhibited a remarkable strand bias toward the C-rich strand (Supplementary Figure S12A). Moreover, the extent of strand bias appeared to correlate with C-richness (Supplementary Figure S12B). The extreme characteristic of the nucleotide composition of C3D lead us to consider that the occurrence of a technical artifact of the TACS-T4 scheme. However, the libraries prepared with two different protocols also showed these features (Supplementary Figure S5 and S12C). Therefore, we concluded that a certain fraction of C3D (i.e., peak-forming C3D) present such characteristic nucleotide composition.

**Figure 7.**
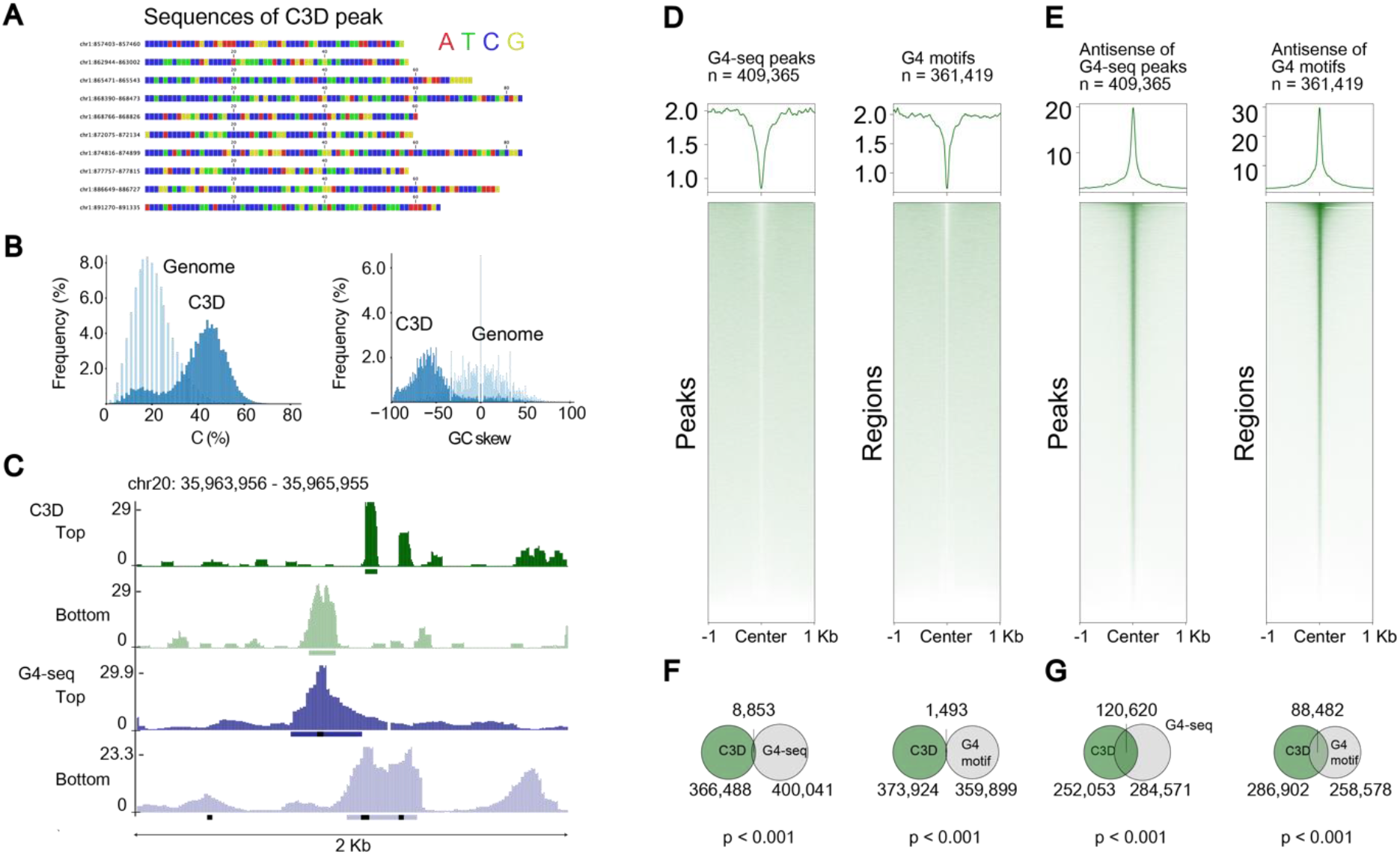
One-third of C3D peaks are complementary strands of G4 motifs. **A.** Nucleotide sequences of representative C3D peaks are displayed using color-coding. **B.** Distribution of C-content (left) and GC-skew (right) are shown for C3D peaks (blue) and 50-mers randomly picked from the reference genome sequence (light blue). **C.** A genome browser shot for C3D and G4-seq (Chambers et al. 2015). The reads mapped to the top and bottom stands are displayed in separate tracks. Horizontal bars at the bottom of each track indicate MACS2-called peaks. The black dots at the bottom of tracks indicate the putative G4 sequences identified using the quadparser algorithm (Huppert and Balasubramanian 2005). **D, E.** Aggregation plots of C3D read coverage on the peaks of G4-seq (left) and the putative G4 sequences identified using the quadparser algorithm (right). The results are shown for the G4-containing strand (D) and its antisense strand (E). **F, G.** Venn diagrams indicating the overlap of C3D peaks with G4-seq peaks (left) and G4-motifs (right). The results are shown for the G4-containing strand (F) and its antisense strand (G).

Since these C3D sequences have several consecutive C tracts, we sought to determine whether C3Ds are rich in the i-motif or the complementary strand of the G-quadruplex (G4) motif. As exemplified in the genome browser shot in Figure 7C, C3D peaks often colocalized with peaks of G4-seq (Chambers et al. 2015). We thus conducted systematic colocalization analyses of C3D reads with both the peaks defined by G4-seq (Chambers et al. 2015) and the G4 motifs predicted with the *quadparser* algorithm (Huppert and Balasubramanian 2005). The sense strands of G4-seq peaks and G4 motifs (i.e., G4-containing strands) failed to enrich, or even excluded, C3D reads (Figure 7D). In contrast, their antisense strands exhibited strong colocalization with C3D reads (Figure 7E). Strikingly, 32.1% and 23.6% of C3D peaks overlapped with the antisense strands of G4-seq peaks and G4 motifs, respectively (Figure 7F and G, Supplementary Figure S13). Intriguingly, C3D^on^ showed a prominent antisense-specific enrichment to a half of G4 motifs but barely overlapped with the other half, whereas C3D^off^ exhibited a much weaker enrichment to almost all G4 motifs (Figure S14A and B). We thus concluded that the genomic origins of C3D frequently overlap with the G4 structure and that the antisense strands of these regions comprise as much as one-third of C3D.

We conducted the same analysis on the data by Snyder et al. (Snyder et al. 2016) and Burnham et al. (Burnham et al. 2016); however, we failed to identify any correlation between their short cfDNA peaks and G4 motifs (Supplementary Figure S14C–F). The discrepancy between our results and theirs likely stems from differences in the cfDNA purification method chosen, since the kit used in their studies cannot quantitatively recover short cfDNA, including C3D (Figure 1C, Supplementary Figures S1D and Supplementary Table S2). Supporting this hypothesis, when we applied their library preparation methods to the cfDNA purified with the PPIP method, we observed enrichments of the C3D reads on both the peaks of the G4-seq (Chambers et al. 2015) and the G4 motifs predicted with the *quadparser* algorithm (Huppert and Balasubramanian 2005) (Supplementary Figure S15).

Finally, we investigated how the G4 structures contributed to the localization specificity of the C3D peaks on the genomic features. We selected C3D peaks without G4 motifs, which comprised 51% of the total C3D peaks (Supplementary Figure S16A), and then subjected them to the same enrichment assay as performed for Figure 5B. Intriguingly, we did not observe enrichment of the promoter, 5′ UTR, and the low complexity region for the peaks without G4 motifs (Supplementary Figure S16B). On the other hand, the enrichment of CpG island and simple repeats was the same regardless of the presence of G4 motifs (Supplementary Figure S16B). These results indicated that at least two types of C3D peaks exist, and these peaks have different structural characteristics.

## DISCUSSION

In the present study, we investigated the cell-free fraction of blood to determine whether, and to what extent, it contains DNA fragments shorter than the well-described nucleosome-sized cfDNA. By combining a conventional method for nucleic acid purification (the PPIP method), to improve the recovery of short fragments (Figure 1A), with a 3′-end labeling method, to improve the detection of short fragments (Figure 1B), we revealed a previously overlooked class of blood cfDNA, termed C3D (Figure 1C–E). C3D is an ssDNA molecule of approximately 50 nt in length (Figure 1C–E) that exists in the blood in an unprotected form at a comparable molar concentration with the NPD (Figure 1D, 1E and Supplementary Figure S2). To determine the nucleotide sequence of C3D, we established a library preparation protocol based on our recently developed, unique ssDNA ligation technique (Figures 2A and B). In-depth sequence analysis of the cfDNA libraries showed that C3D was derived from open chromatin regions (Figure 6) and transcription factor-binding sites (Figure 6 and Supplementary Figure S9). Moreover, as much as one-third of C3D peaks corresponded to the antisense strand of putative G-quadruplex structures (Figure 7). Based on these previously undescribed features, we propose that C3D is a novel class of plasma cfDNA that long escaped detection as it is not quantitatively recovered by the popular cfDNA isolation method (Figure 1C and Supplementary Figure S1D) and cannot be converted to sequenceable forms unless ssDNA-compatible protocols are used.

The discovery of C3D has raised several new questions to be addressed in future studies, including the mechanism(s) leading to its production. cfDNA is believed to be generated by nuclease digestion because of apoptotic cell death (Aucamp et al. 2018; Watanabe et al. 2019; Han et al. 2020). Moreover, while NPD is double-stranded, C3D is single-stranded (Figure 1D and E). Why is C3D single-stranded? If the genomic origins of C3D conform to the canonical dsDNA structure, then how and when are they converted to single-stranded forms? The important facts required to address these questions are that the base composition of C3D is strongly biased toward C-richness (Figure 7B and Supplementary Figure S12) and that one-third of C3D peaks are the complementary strands of G4 structures, which may form the i-motif structure (Abou Assi et al. 2018) (Figure 7E and G). G4 structures are enriched in the regulatory regions of genes such as promoters and nucleosome-free regions (Huppert and Balasubramanian 2007; Hansel-Hertsch et al. 2016). The enrichment of C3D peaks in promoters, CpG islands (Figure 5), DHS, and ATAC-Seq peaks (Figure 6) could be partially explained by this G4 abundance of antisense C3D. In this sense, during the generation of the G4 structure its complementary strand is released, which in part explains the mechanism of C3D production. Alternatively, the formation of the i-motif structure generates an unstructured anti-sense G-rich strand, as the formation of the i-motif and G4 structures are mutually exclusive. When released in the plasma, the structured i-motif is conceivably more stable than its unstructured complementary strand, thus leading to the observed enrichment of C-rich strands in C3D. To fully address these queries, it is critical to understand whether the production of C3D can be recapitulated in model animals and cell lines.

Another question is whether C3D reflects any pathophysiological conditions, including sex, age, circadian rhythm, pregnancy, and various diseases, like the NPD or conventional cfDNA (Lo et al. 2010; Burnham et al. 2017; Cohen et al. 2017; Bronkhorst et al. 2019). Herein, we have shown that C3D and NPD are derived from distinct genomic regions (Figures 5C and D). Hence, they may reflect different pathophysiological conditions. It is worth investigating whether the amount and composition of C3D alter depending on the health conditions of the donors. Our preliminary data suggest that C3D differs between healthy individuals and patients with cancer (data not shown). It is also intriguing to examine whether C3D is present in body fluids other than blood, such as urine and cerebrospinal fluid.

Further investigations are required to adequately explore the biology and potential applications of C3D. To facilitate these investigations, it is necessary to improve C3D sequencing using the TACS-T4 method. The most important issue is the suppression of adapter dimer formation. In the second adapter tagging, T4 DNA ligase produces adapter dimers from two phosphorylated adapters. Since the sizes of the dimer and the library molecule are similar, selective removal of the former is not easy, necessitating labor-intensive, time-consuming steps. Thus, a method that does not include formation of adapter dimers would be required for a more sensitive and practical library preparation from C3D. It is also desirable to develop a simple, multiplexable method to isolate cfDNA, including C3D, from various body fluids. These, and other techniques, would accelerate the exploration of C3D.

## METHODS

### Blood samples

The ethics review board at Kyushu University approved the procedure for collecting blood samples and their use for genome sequencing (approval I.D. 752-00). We used anonymized blood samples collected from five healthy males after obtaining their written informed consent. For the isolation of plasma, blood was drawn into BD vacutainer EDTA-2K collection tubes (Becton Dickinson, Franklin Lakes, NJ, USA) and centrifuged at 1,300 ×*g* for 10 min at 4 °C. For serum separation, blood was drawn into a BD vacutainer plain tube (Becton Dickinson), incubated at room temperature for 30 min, and centrifuged at 1,300 ×*g* for 10 min at 4 °C. The plasma and serum were again centrifuged at 14,000 ×*g* for 10 min at 4 °C to minimize the contamination of cellular DNA. Plasma and serum separations were performed within 30 min of blood collection. After obtention, plasma and serum samples were immediately stored at −20 °C until use.

Plasma and serum of healthy individuals were also obtained from BIOPREDIC (Rennes, France), Cosmo Bio (Tokyo, Japan), and Clinical Trials Laboratory Services (London, UK). The details for the blood samples used in this study are summarized in Supplementary Table S6.

### Purification of cfDNA

For most of this study, cfDNA was prepared using the PPIP method as follows: Plasma/serum (500 μL) was combined with 12 μL of 5 M NaCl, 10 μL of 500 mM ethylenediaminetetraacetic acid (EDTA), 30 μL of 10% (w/v) sodium dodecyl sulfate (SDS), and 10 μL of 20 mg/mL proteinase K (Qiagen) and incubated at 60 °C for 30 min. Protease-treated plasma/serum was extracted with 600 μL of phenol, 600 μL of phenol-chloroform, and 600 μL of chloroform. The aqueous phase was transferred to a new tube, combined with 60 μL of 3 M sodium acetate (pH 5.2) and 660 μL of isopropanol, and subjected to centrifugation at 20,000 ×*g* for 10 min. The DNA pellet was rinsed with 70% (v/v) ethanol and dissolved in 5–10 μL of 10 mM Tris-HCl (pH 8.0).

In certain parts of this study, the cfDNA was isolated using commercially available kits in order to compare them with the PPIP method. These kits included QIAamp Circulating Nucleic Acid Kit (Qiagen, Hilden, Germany), Plasma/Serum Cell-Free Circulating DNA Purification Mini Kit (Norgen Biotek, Thorold, Canada), NucleoSpin Plasma XS (Takara Bio Inc., Shiga, Japan), and NEXTprep-Mag cfDNA Isolation Kit (PerkinElmer, Waltham, MA, USA), according to the manufacturer's instructions. The name of the kits used to obtain the data of a specific figure is indicated in the pertinent figure legend.

DNA concentration was measured with the Qubit ssDNA Assay Kit and Qubit dsDNA HS Assay kit on a Qubit Fluorometer (Thermo Fisher Scientific, Waltham, MA, USA). The purified DNA was stored at −20 °C until use.

### Analysis of cfDNA with denaturing polyacrylamide gel electrophoresis

The following were added to 4 μL of purified cfDNA: 4 μL of 2.5× TACS buffer (125 mM HEPES-KOH (pH 7.5), 12.5 mM MgCl_2_, 1.25% (v/v) Triton-X100, and 50% (w/v) polyethylene glycol 6000 [Nacalai Tesque, Kyoto, Japan]), 1 μL of 0.05 mM 7-propargylamino-7-deaza-ddATP-6-FAM (Jena Bioscience, Jena, Germany), 0.4 μL of internal standard solution (see Supplementary Methods), and 0.5 μL of TdT (Takara Bio Inc.); the reaction volume was adjusted to 15 μL with water. After incubating the reaction mixture at 37 °C for 30 min, 7.5 μL of buffer B2 (3 M guanidine hydrochloride, 20% (v/v) Tween 20), and 1 μL of 20 mg/mL proteinase K were added, and the mixture was incubated at 55 °C for 15 min. Next, DNA was recovered with solid-phase reversible immobilization (SPRI) (DeAngelis et al. 1995) as follows: The proteinase K-treated reaction was combined with 1 μL of Sera-Mag carboxylate-modified magnetic particles (GE Healthcare, Chicago, IL, USA.) and 72 μL of binding buffer (300 mM NaCl, 3 mM Tris-HCl [pH8.0], 0.3 mM EDTA, 0.015% (v/v) Tween 20, 70% (v/v) ethanol). After incubation at room temperature for 5 min, the beads were washed with 70% (v/v) ethanol. Purified DNA was eluted with 4 μL of 10 mM Tris-acetate (pH 8.0) and analyzed on a 10% Novex TBE-Urea gel (Thermo Fisher Scientific). Fluorescent images were obtained using Typhoon Trio+ and analyzed with ImageQuant software (GE Healthcare).

### Analysis of the plasma ultracentrifugation fraction

Plasma and serum were first centrifuged at 10,000 ×*g* for 30 min at 4 °C, and the supernatant was used for ultracentrifugation. One milliliter of the precleared plasma or serum was transferred to a thick-wall polypropylene tube (Beckman Coulter, Brea, CA, USA) and centrifuged in an OptimaMAX (Beckman Coulter) benchtop ultracentrifuge with a TLS-55 rotor for 70 min at 100,000 ×*g* at 4 °C. The supernatant was saved for cfDNA purification and western blotting without any further preparation. The pellet was resuspended in 1 mL of phosphate buffered saline (PBS) and centrifuged again for 70 min at 100,000 ×*g* at 4 °C. After removing the supernatant, the pellet was dissolved in 50 μL of 1× SDS sample buffer, divided into two portions, and saved for DNA analysis and western blotting. Western blotting was performed using a monoclonal antibody raised against CD9 (catalog number 014-27763, FUJIFILM Wako Chemicals, Osaka, Japan). For details, see the Supplementary Methods.

### Production of recombinant enzymes

For TACS ligation, we used TS2126 RNA ligase prepared in-house with certain modifications to the previously described method (Miura et al. 2019). For details, see the Supplementary Methods. CircLigase II can be used instead of TS2126 RNA ligase.

Recombinant human aprataxin was also prepared in-house. DNA fragment encoding human aprataxin (UniProt#Q7Z2E3) was chemically synthesized by Eurofins Genomics (Tokyo, Japan) with codon optimization for *E. coli* (Supplementary Information) and subcloned into pColdI (Takara Bio Inc.) for protein expression in *E. coli*. For details, see the Supplementary Methods. The expression vector is available from the authors upon request; however, only to those who have an appropriate license to use pColdI.

### Library preparation from cfDNA based on TACS ligation (TACS-T4 scheme)

First, 10 ng of cfDNA was dephosphorylated in a 10-μL reaction containing 2.5 μL of 10× TACS buffer and 1 μL of shrimp alkaline phosphatase (Takara Bio Inc.) at 37 °C for 15 min. The reaction mixture was then heated at 95 °C for 5 min to inactivate the enzyme and denature the DNA. Next, adapter tagging of single-stranded cfDNA was performed with TACS ligation (Miura et al. 2019). The 10 μL reaction mixture after dephosphorylation was supplemented with 10 μL of 50% (w/v) PEG, 1 μL of 10 μM PA-TruSeqIndex-dSp-P (Supplementary Table S7), 1 μL of 10 mM ATP, 1 μL of TdT (Takara Bio Inc.), and 1 μL of 2 mg/mL TS 2126 RNA ligase (Supplementary Methods). The reaction mixture was then sequentially incubated at 37 °C for 30 min, 65 °C for 2 h, and 95 °C for 5 min. Next, DNA complementary to the adaptor-tagged DNA was synthesized. After adapter tagging, 5 μL of 10× ExTaq buffer (Takara Bio Inc.), 5 μL of 2.5 mM dNTPs (Takara Bio Inc.), 1 μL of 20 μM TruSeqUniv (Supplementary Table S7), 1 μL of 2.5 U/μL hot-start Gene Taq (Nippon Gene), and 1 mg/mL aprataxin, were added. The total volume was adjusted to 50 μL with water. The reaction mixture was then sequentially incubated at 37 °C for 15 min, 95 °C for 3 min, 55 °C for 5 min, and 72 °C for 5 min. Subsequently, 1 μL of T4 DNA ligase (Takara Bio Inc.) was added, and the reaction mixture was incubated at 25 °C for 1 h. Finally, the library DNA was purified using SPRI. After the second adapter ligation, 25 μL of buffer B2 and 5 μL of 20 mg/mL proteinase K were added. After incubation at 50 °C for 15 min, the reaction mixture was combined with 146 μL of AMPure XP (Beckman Coulter) and incubated at room temperature for 5 min to capture the DNA on the surface of the beads. The beads were collected using a magnet and rinsed with 70% ethanol, and the library DNA was eluted in 25 μL of 10 mM Tris-acetate (pH 8.0).

The library was amplified by PCR for the completion of library molecule structure and indexing. To the 25 μL elute, 25 μL of 2× PrimeStar Max, 0.4 μL of PCR-Univ, and 0.4 μL of PCR-Index primer (see Supplementary Tables S7 and S8), were added. Following incubation at 95 °C for 1 min, the reaction mixture was subjected to 10 cycles of 3-step incubations at 95 °C for 10 s, 55 °C for 15 s, and 72 °C for 30 s. Next, 50 μL of the amplified library was combined with 75 μL of AMPure XP, and the suspension was incubated at room temperature for 5 min. Beads were collected and rinsed with 70% (v/v) ethanol, and DNA was eluted with 50 μL of 10 mM Tris-acetate (pH 8.0). This SPRI-based purification method was repeated five times to remove the adapter dimer. The molar concentration of the library was determined by quantitative PCR (qPCR) using Library Quantification kit (Takara Bio Inc.) according to the manufacturer’s instructions. The amplified PCR product was analyzed by denaturing gel electrophoresis using 6% Novex TBE-Urea gel.

### Other library preparation methods

The dsDNA ligation-based method was also compared. The ThruPLEX DNA-Seq Kit (Takara Bio) was used following the manufacturer’s instructions.

Methods based on two different principles for ssDNA ligation were also compared with the TACS-T4 scheme. The first one was the CircLigase II-based method (Gansauge and Meyer 2013), and the other was based on T4 DNA ligase (Gansauge et al. 2017). We faithfully followed the original protocols described in the literature except for the PCR amplification steps, in which PrimeStar Max was used as described above.

### Sequencing

Small-scale sequencing was performed using Illumina MiSeq with MiSeq Reagent Kit v3 (150 cycles) in the paired-end mode of 2× 75 cycles. For large-scale sequencing, paired-end sequencing with 2× 150 cycles using the HiSeq X Ten was performed by Macrogen Japan Corp. (Kyoto, Japan). The reads were delivered after demultiplexing, and indexed libraries were used for subsequent bioinformatics analysis.

### Bioinformatic analysis

Sequenced reads were first filtered using fastp (Chen et al. 2018), and the additional nucleobases attached during TACS-T4 library preparation were trimmed from both ends of the reads using SeqKit subseq (Shen et al. 2016). The processed reads were then mapped to the reference human genome assembly GRCh37 (hg19) with Bowtie2 in paired-end mode (Langmead and Salzberg 2012). The alignments uniquely mapped to the genome were separated based on their fragment size into C3D or NPD; fragments ranging from 35 to 75 nt were defined as C3D, whereas those ranging from 147 to 190 nt were defined as NPD. The alignments mapped to the top and bottom strands of the reference genome were then divided and individually subjected to peak calling with MACS2 (Zhang et al. 2008). Next, the MACS2-called peaks were merged into a single file. The strand-specific BAM files were also converted to BigWig format using BEDTools genomecov (Quinlan and Hall 2010) and visualized using the UCSC genome browser with Trackhub function (Kent et al. 2002; Raney et al. 2014). The coverage of mapped cfDNA fragments around the human genes or known genomic regions was plotted using the deepTools computematrix and plotHeatmap (Ramirez et al. 2016). HOMER annotatePeaks were used to link the peaks with known functional elements (Heinz et al. 2010). ChIPPeakAnno (Zhu et al. 2010) was used to construct Venn diagrams of overlapping peaks, and regioneR (Gel et al. 2016) was used for the permutation test. The sequences of the C3D peaks were extracted using the getFasta of BEDTools (Quinlan and Hall 2010). The base composition of each DNA sequence was calculated with fx2tab of SeqKit (Shen et al. 2016) and visualized with ggplot2 (Wickham 2016). The flowcharts for analytical pipelines are provided in Supplementary Information S3.

## DATA ACCESS

The sequence data used in the present study were deposited in the Japanese Genotype-Phenotype Archive (JGA) under the accession number JGAS000257. The publicly available data used in the current study is summarized in Supplementary Tables S9 and S10.

## ACKNOWLEDGMENTS

We are grateful to Goro Doi, Miki Miura, and Yukiko Shibata for their assistance in collecting blood samples and preparing enzymes. We also appreciate the technical assistance from the Research Support Center, Research Center for Human Disease Modeling, Kyushu University Graduate School of Medical Sciences; Department of Clinical Chemistry and Laboratory Medicine, Kyushu University Graduate School of Medical Sciences; and the Cooperative Research Project Program of the Medical Institute of Bioregulation, Kyushu University. We would like to thank Editage (http://www.editage.com) for English language editing.

