## Supplementary Information for "Short single-stranded DNA with putative non-canonical structures comprises a novel class of plasma cell-free DNA"

#### Supplementary Figures

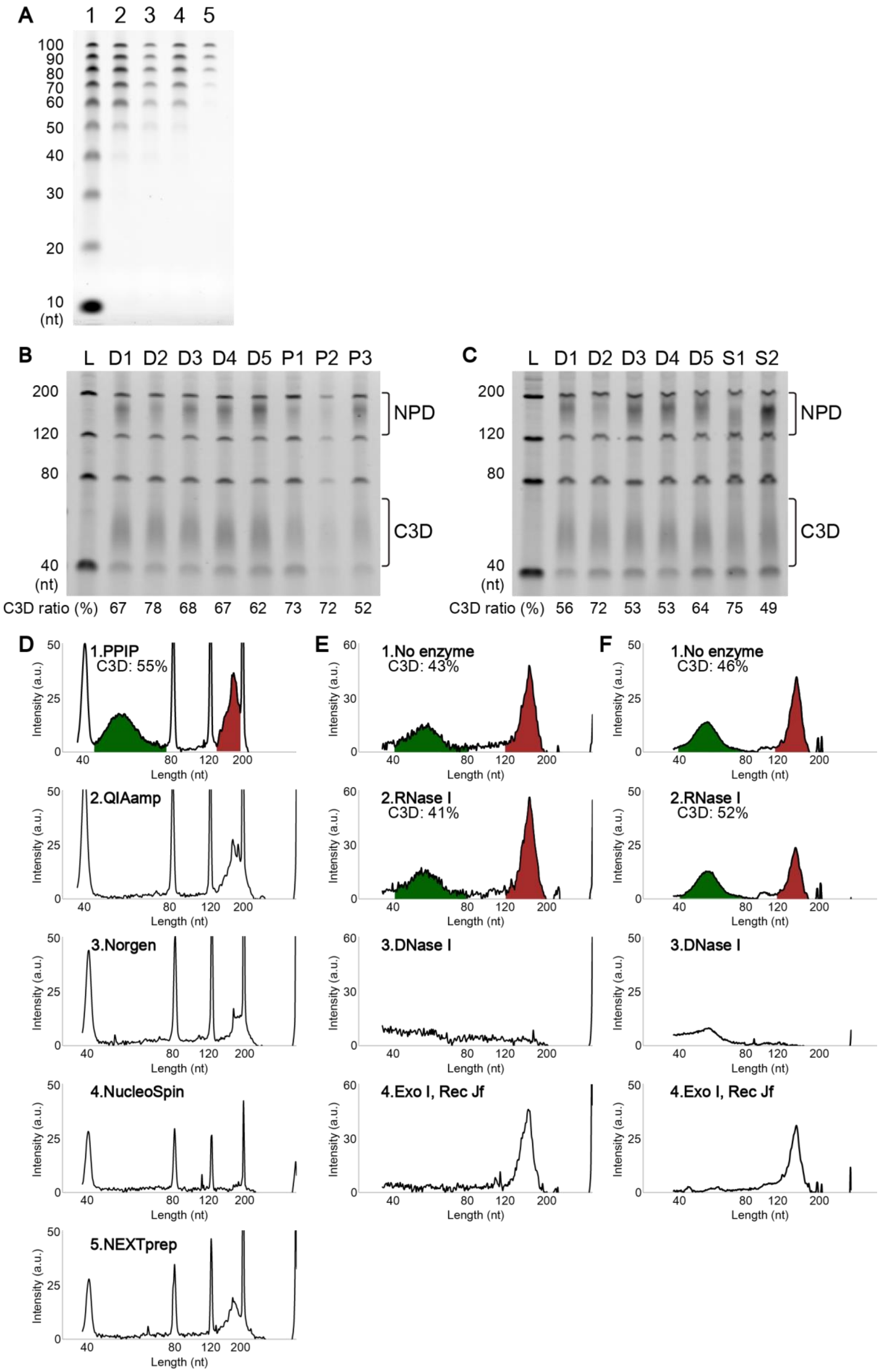


*(Continues on the next page)*

**Supplementary Figure S1. Short single-stranded DNA in cell-free blood fractions.**

A. DNA samples recovered using four commercially available kits were compared. As a model, the 10 bp DNA step ladder from Promega (Fitchburg, WI, USA) was used (lane 1). The kits used were the QIAamp Circulating Nucleic Acid Kit (Qiagen, Hilden, Germany) (lane 2), the Plasma/Serum Cell-Free Circulating DNA Purification Mini Kit from Norgen Biotek (Thorold, Canada) (lane 3), the NucleoSpin Plasma XS from Takara Bio Inc. (Shiga, Japan) (lane 4), and the NEXTprep-Mag cfDNA Isolation Kit from PerkinElmer (Waltham, MA, USA) (lane 5).

B and C. The cfDNAs purified from plasma (B) and serum (C) were compared. Plasma and serum were obtained from five healthy donors and commercial sources (see Supplementary Table S6). For details of the experimental procedures, see Supplementary Methods.

D–F. The electropherograms of the gel images from Figure 1C (D), Figure 1D (E), and Figure 1E (F). The areas for C3D (green, 40–80 nt) and NPD (red, 120–200 nt) were calculated, and the relative intensity of C3D is indicated. a.u.; arbitrary unit.

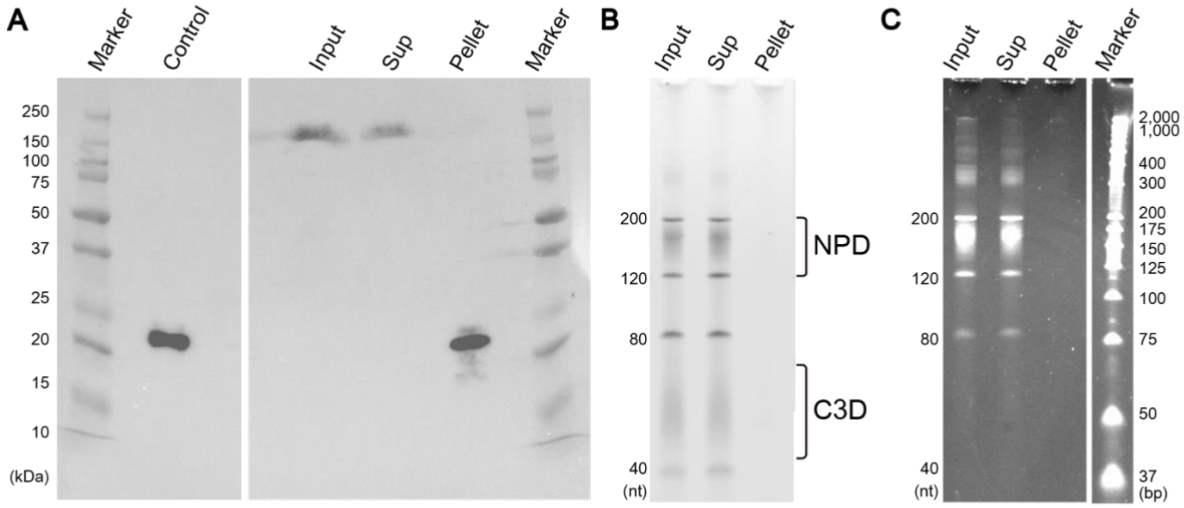


**Supplementary Figure S2. The membranous fraction of the plasma does not contain cfDNA.**

The plasma from a healthy individual (Donor 1) was fractionated via ultracentrifugation. The supernatant and pellet were used for western blotting and cfDNA purification.

A. Western blotting was performed using an antibody raised against the exosome specific protein marker CD9. As a positive control, purified exosomes from COLO201 cells (FUJIFILM Wako, Osaka, Japan) were loaded (Control). The loading amounts of the plasma before ultracentrifugation (Input) and the supernatant after separation (Sup) were equivalent, whereas 20 times the equivalent amount was loaded for the pellet (Pellet).

B. The cfDNAs purified from the plasma (Input), supernatant (Sup), and pellet (Pellet) after ultracentrifugation were analyzed by denaturing gel electrophoresis. As an internal control, the oligonucleotide mixture (see Supplementary Methods) was spiked into the plasma before centrifugation. After purification, cfDNA was fluorescently labeled and loaded onto the gel. As in A, the input (Input) and the supernatant (Sup) loading amounts were equivalent, whereas 20 times the equivalent amount was loaded for the pellet (Pellet).

C. The gel analyzed in B was stained with SYBR Gold nucleic acid gel stain (ThermoFisher Scientific, Waltham, MA, USA).

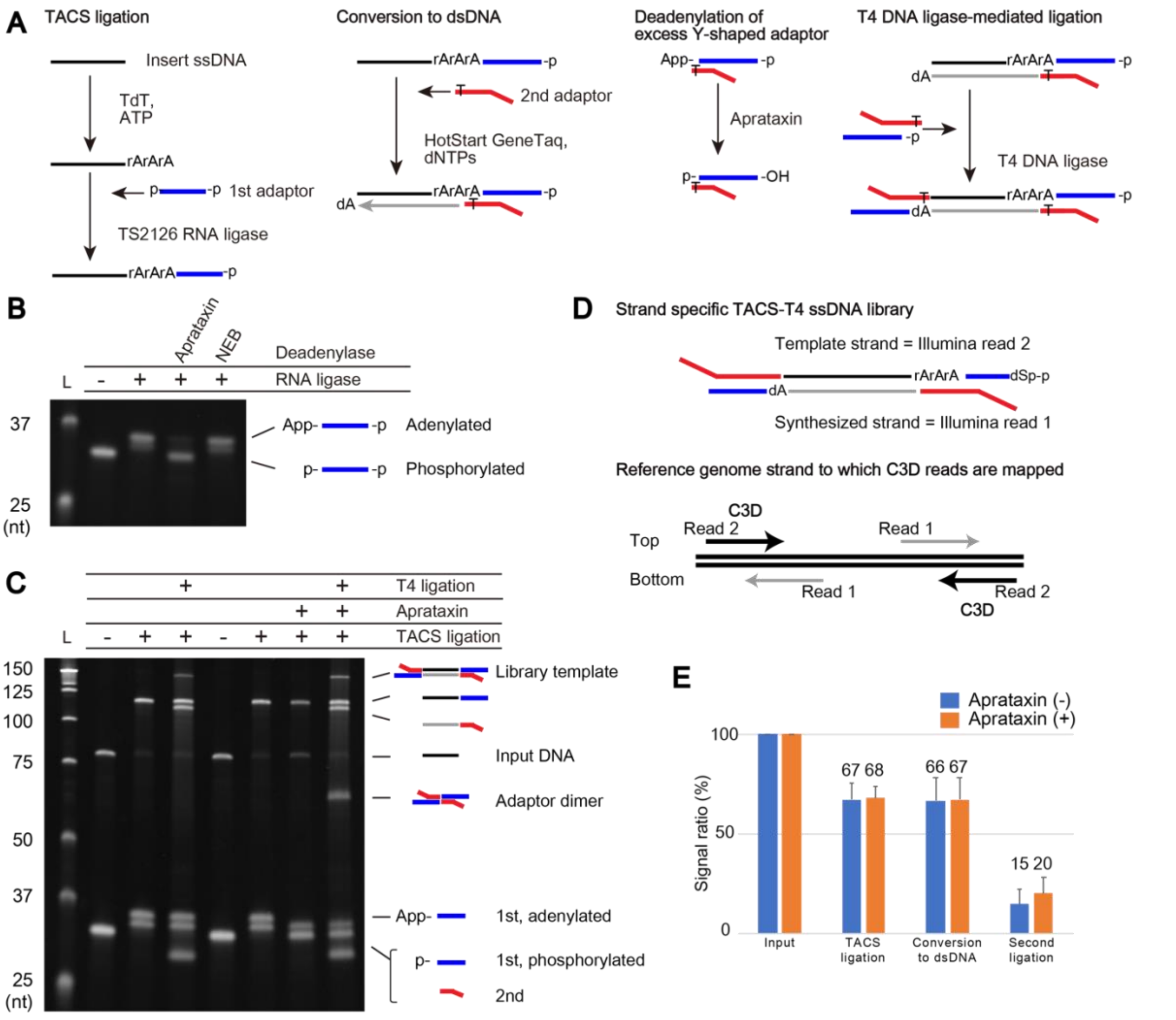


**Supplementary Figure S3. TACS-T4 scheme for the library preparation from ssDNA.**

A. The details of each step of the TACS-T4 scheme.

B. Aprataxin-mediated deadenylation of the adaptor. The TS2126 RNA ligase used in TACS ligation adenylates the 5′-phosphorylated end of the adaptor. Since the 3′-end of the adaptor is also phosphorylated to repress its concatenation (see Supplementary Table S7 for the structure of the oligonucleotide used) and the RNA ligase also adenylates the 3′-phosphorylated end of the adaptor, two bands are seen after the RNA ligase reaction. It seems that the 5′ adenylation inhibits T4 DNA ligase-mediated ligation. Therefore, an efficient method to remove the adenylates was required. We found that recombinant human aprataxin is suitable for removing the adenylates in the solution following TACS ligation. Although 5' deadenylase is available from New England Biolabs (NEB; Ipswich, MA, USA), this enzyme seems to remove adenylates in the solution following TACS ligation poorly.

C. The efficiencies of adaptor tagging with and without aprataxin were compared. The enzyme enhances the tagging of the second adaptor with T4 ligase.

D. The strand specificity of the TACS-T4 scheme. As the TACS ligation product contains a few adenylates, if a DNA polymerase without reverse transcriptase activity is used, PCR amplification will occur only from the strand without the adenylates. Based on the design of the adaptor, read one and read two correspond to the reverse strand and the forward strand of target DNA, respectively.

E. The amounts of products from each step in the TACS-T4 scheme were calculated based on electrophoresis signals relative to the input DNA signal. Averages and standard deviations of three replicates are shown. A representative image is shown in Supplementary Figure S3C.

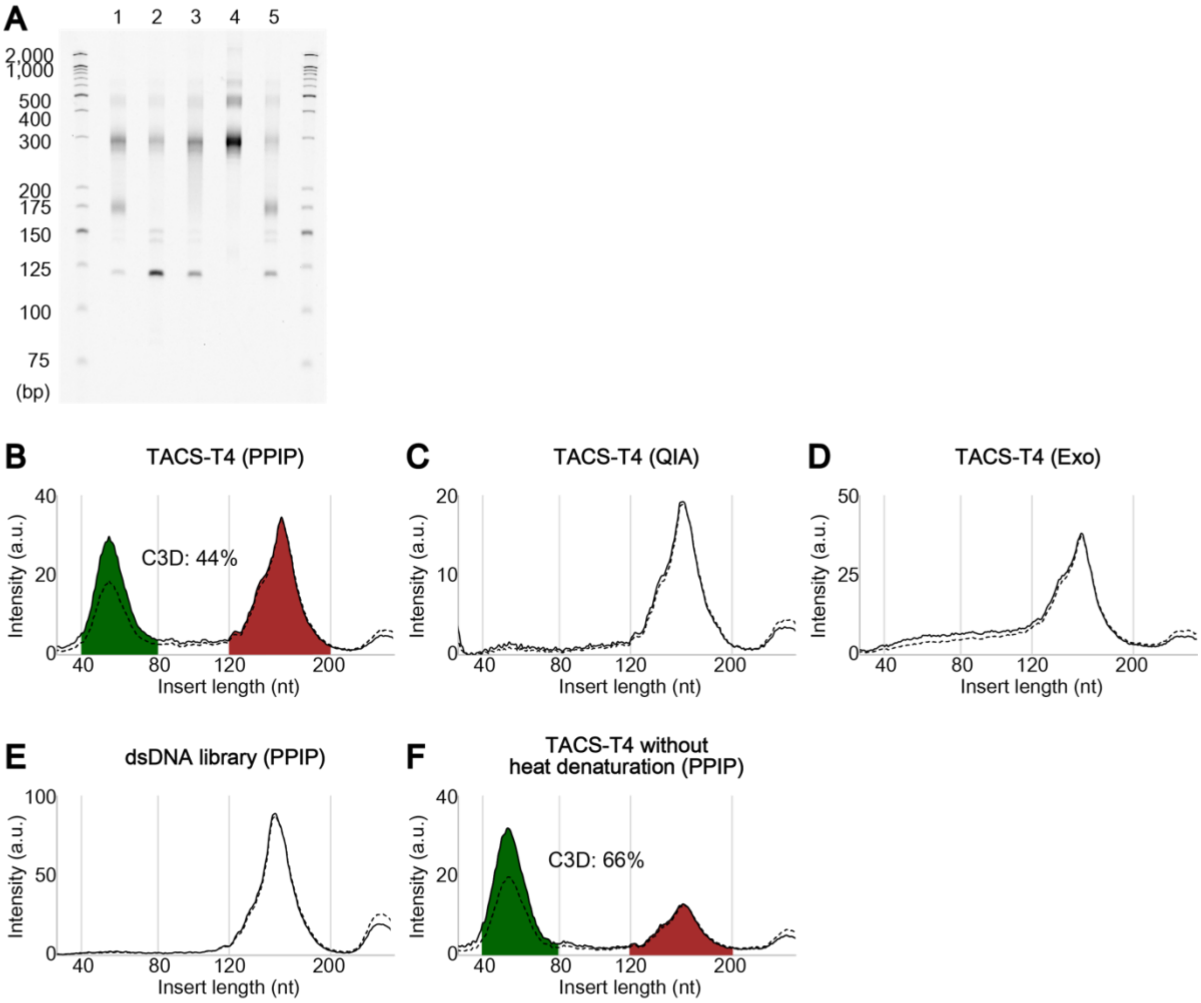


**Supplementary Figure S4. The size distributions of cfDNA fragments in the libraries prepared with different pretreatments reflect different forms of DNA.**

A. The gel image from denaturing gel electrophoresis of amplified libraries with real-time PCR quantification is shown. The gel was stained with SYBR Gold nucleic acid gel stain (ThermoFisher Scientific). The loaded libraries were prepared from cfDNA purified using PPIP (lanes 1, 3, 4, and 5) or QIAamp Circulating Nucleic Acid Kit (Qiagen) (lane 2). For lane 2, exonuclease treatment was performed before the cfDNA purification. The libraries were prepared using the TACS-T4 scheme (lanes 1, 2, 3, and 5) or ThruPLEX DNA-Seq kit (Takara Bio Inc.) (lane 4). For lanes 1–3, the cfDNAs were heat-denatured before the TACS-T4 scheme. Note that the libraries shown here are the same as the ones shown in Figure 2C–G.

B–F. The electropherograms of lanes 1 (B), 2 (C), 3 (D), 4 (E), and 5 (F) of the gel image in A. The signal intensity of the raw data (dashed lines) and after normalization with the molecular weight of the DNA (solid lines) are shown. The areas of C3D (green, 40–80 nt) and NPD (red, 120–200 nt) were calculated, and the relative fraction of C3D is indicated for B–F. a.u.; arbitrary unit.

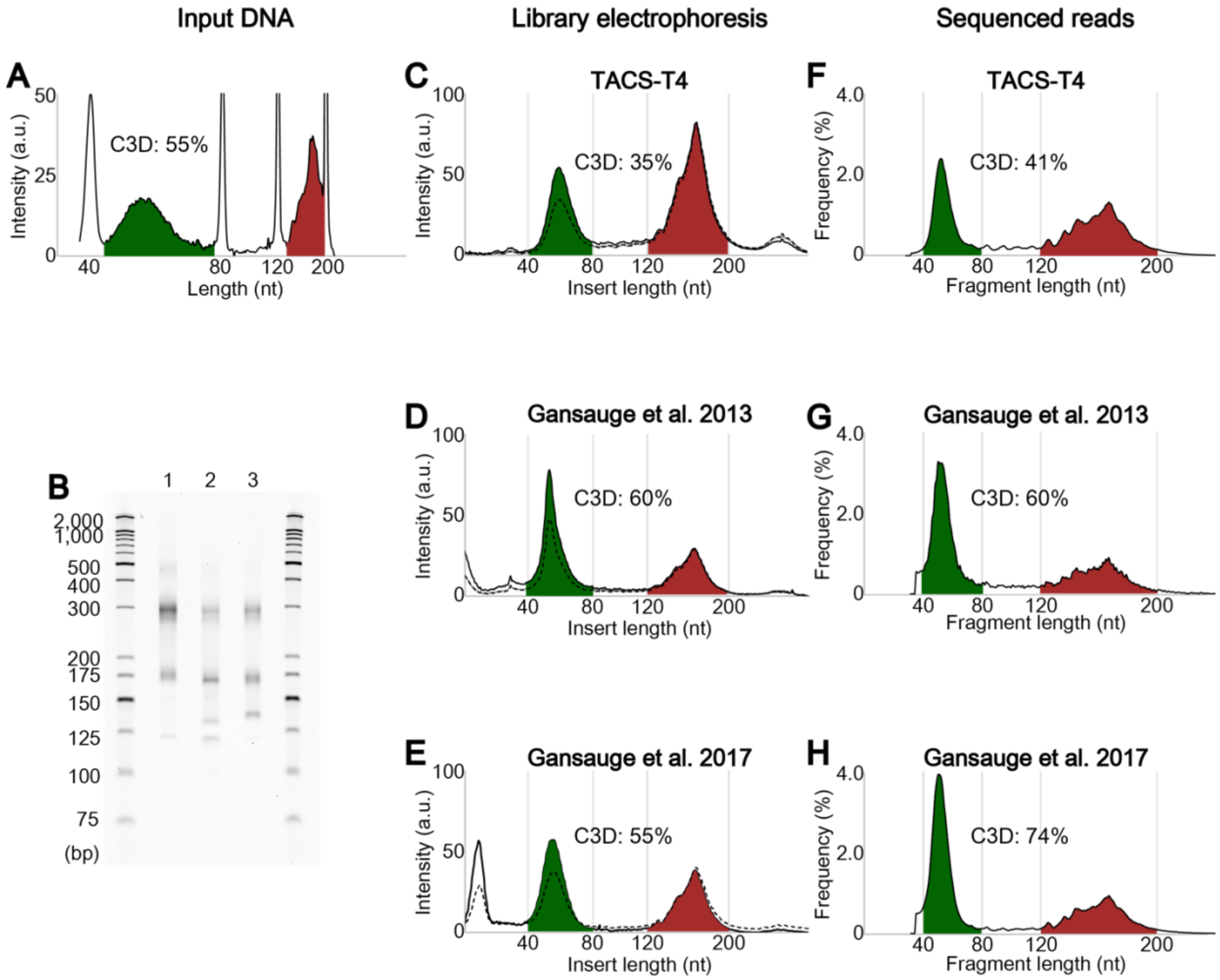


**Supplementary Figure S5. Two major clusters of cfDNA were consistently detected with the three ssDNA-adapted library preparation methods.**

A. The electropherogram of input cfDNA purified with the PPIP method, same as Supplementary Figure S1D.

B. The gel image from denaturing gel electrophoresis of amplified libraries after real-time PCR quantification is shown. The libraries were prepared using the TACS-T4 scheme (lane 1), ssDNA-adapted protocol by Gansauge et al. (2013) (Gansauge and Meyer 2013) (lane 2), or Gansauge et al. (2017) (Gansauge et al. 2017) (lane 3). The gel was stained with SYBR Gold nucleic acid gel stain (ThermoFisher Scientific).

C–E. The electropherograms of lanes 1 (C), 2 (D), and 3 (E) of the gel image in B. The signal intensity of the raw data (dashed lines) and after normalization with the molecular weight of the DNA (solid lines) are shown.

F–H. The size distribution of sequenced reads for libraries, same as lanes 1 (F), 2 (G), and 3 (H) are shown.

The areas of C3D (green, 40–80 nt) and NPD (red, 120–200 nt) were calculated, and the relative fraction of C3D is indicated for C–H. a.u.; arbitrary unit

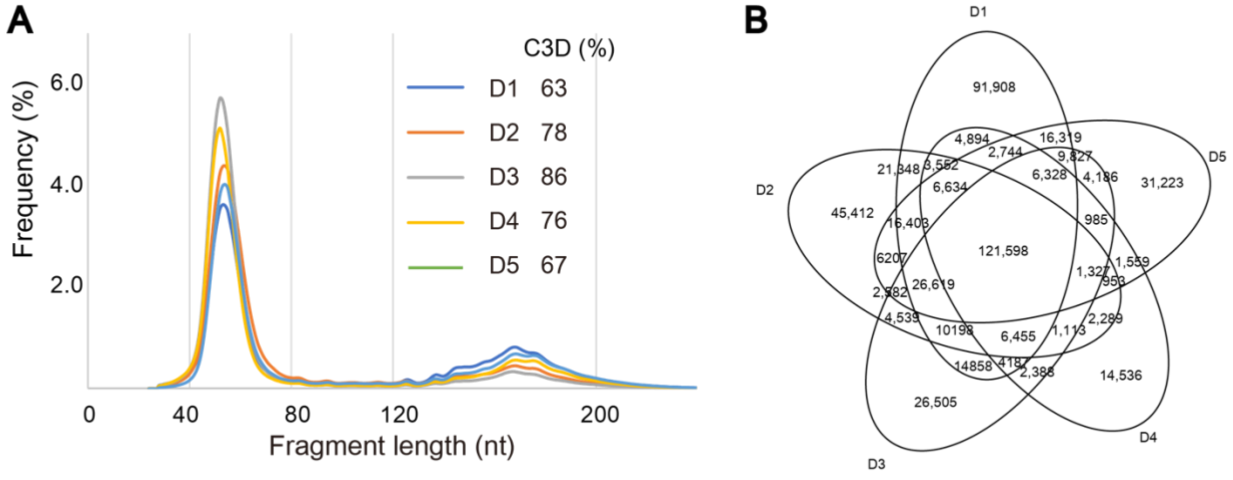


**Supplementary Figure S6. Locations of the C3D peaks are well conserved among individuals.**

A. The size distributions of cfDNA fragments detected by sequencing for five individuals are overlaid. The relative amount of C3D reads are calculated and indicated for each sample.

B. Venn diagram showing overlaps of C3D peaks among the five donors.

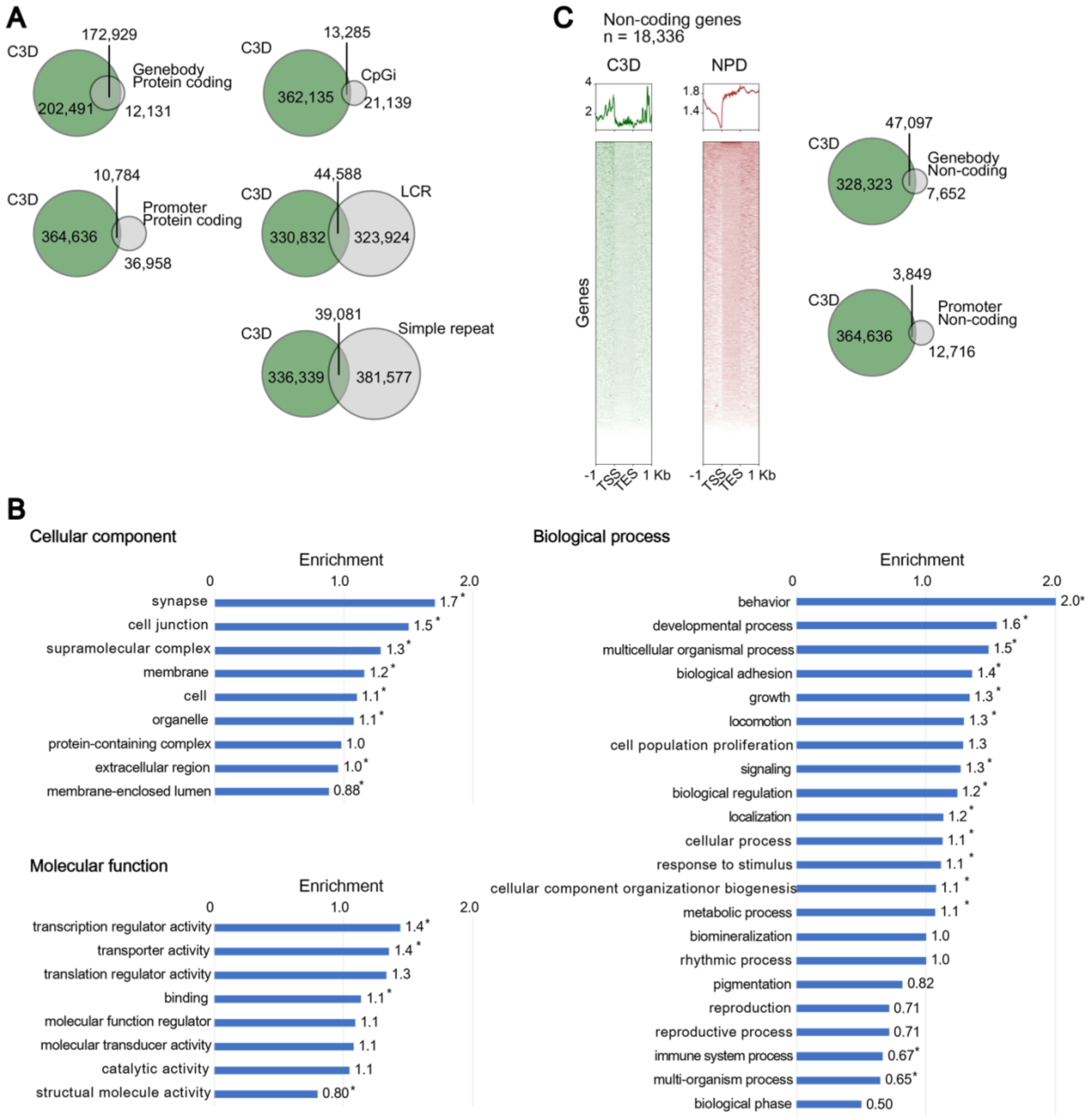


*(Continues on the next page)*

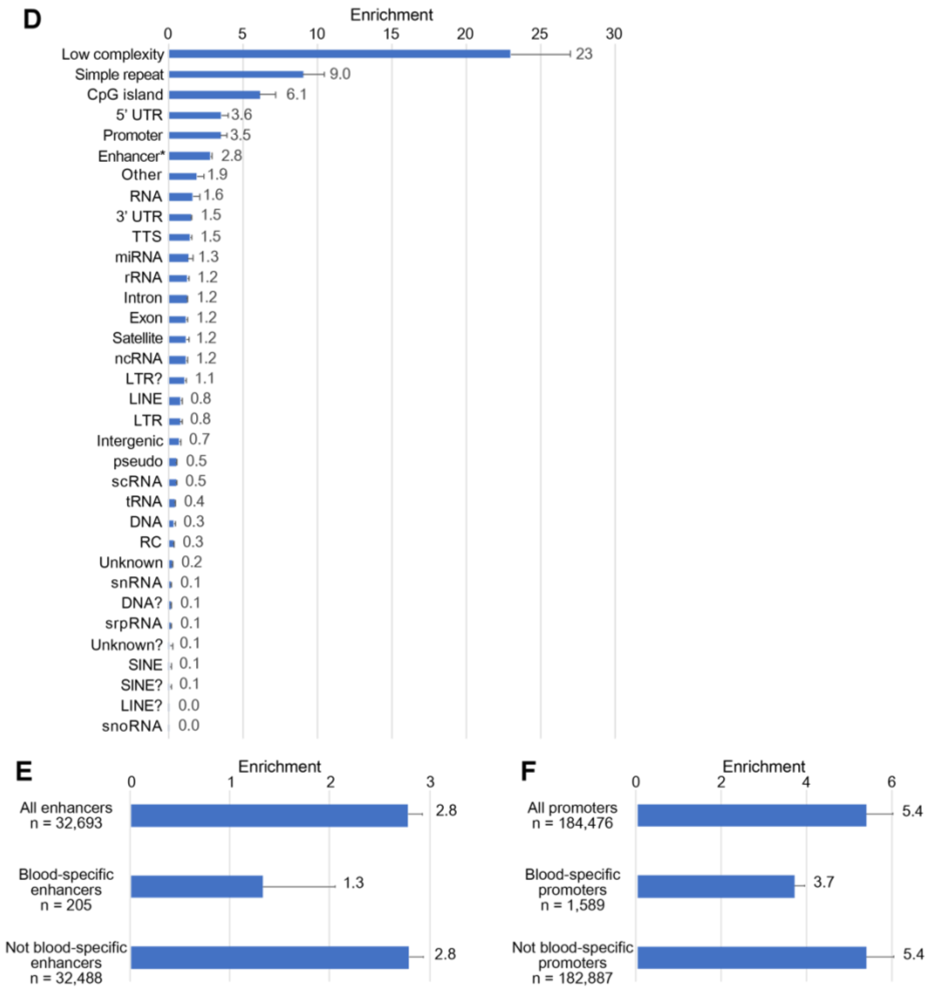


**Supplementary Figure S7. C3D is enriched in the regulatory regions of genes.**

A. The colocalization of C3D peaks and the annotations enriched in the C3D peaks are shown in Venn diagrams.

B. Fold enrichment of gene ontology (GO) terms for genes that contain C3D peaks in their promoters or 5′ UTRs (7,634) compared to that for all reference genes (n = 20,851). *p-value < 0.05 (Fisher's exact test).

C. Aggregation plots of the normalized read coverages of C3D and NPD around the non-coding genes are shown.

D. The enrichment of C3D peaks was calculated for each annotation provided by the "Detailed Annotation" of HOMER annotatePeaks. For enhancers, the data downloaded from the FANTOM5 project (Andersson et al. 2014; Lizio et al. 2019) was used.

E and F. The blood specificity from the enriched C3D peaks in enhancers (E) and promoters (F) was evaluated using the dataset downloaded from the FANTOM5 project (Andersson et al. 2014; Lizio et al. 2019). The blood specificity of enhancers and promoters was defined when the CAGE signal detected for the blood cells exceeded 90% of the total CAGE signal assigned.

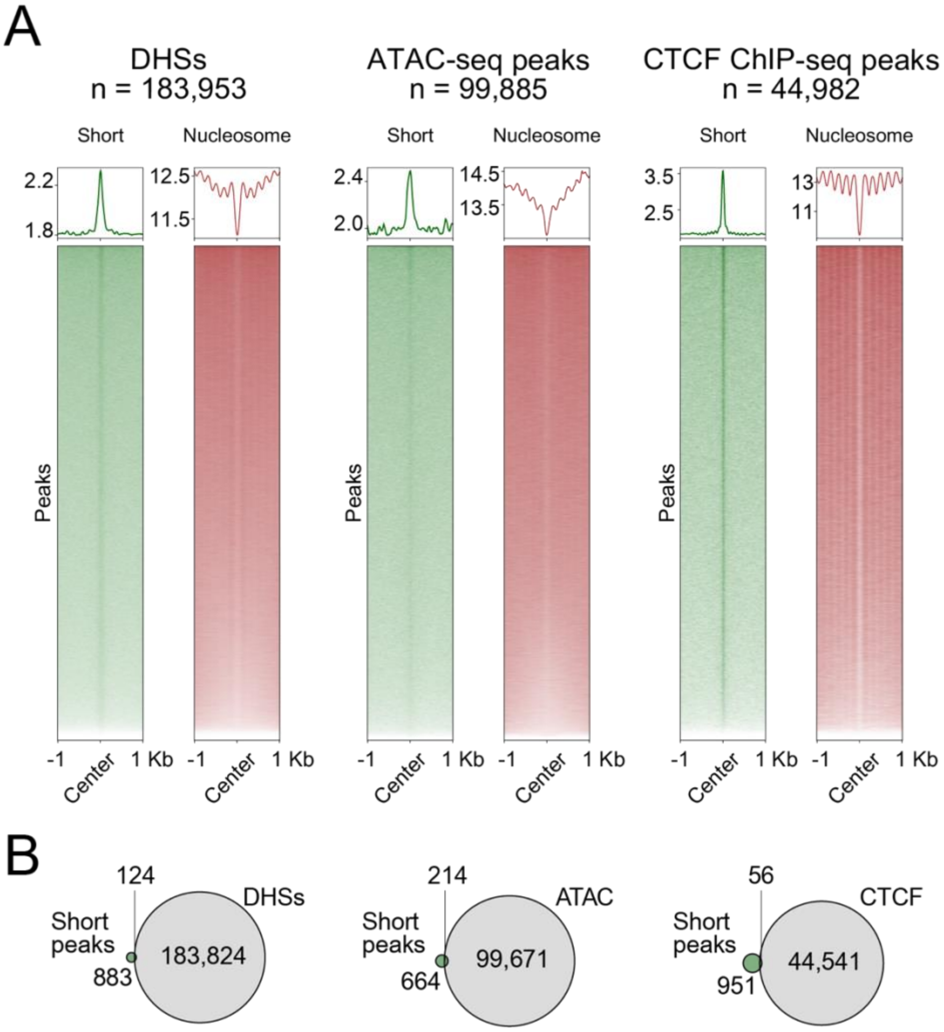


*(Continues on the next page)*

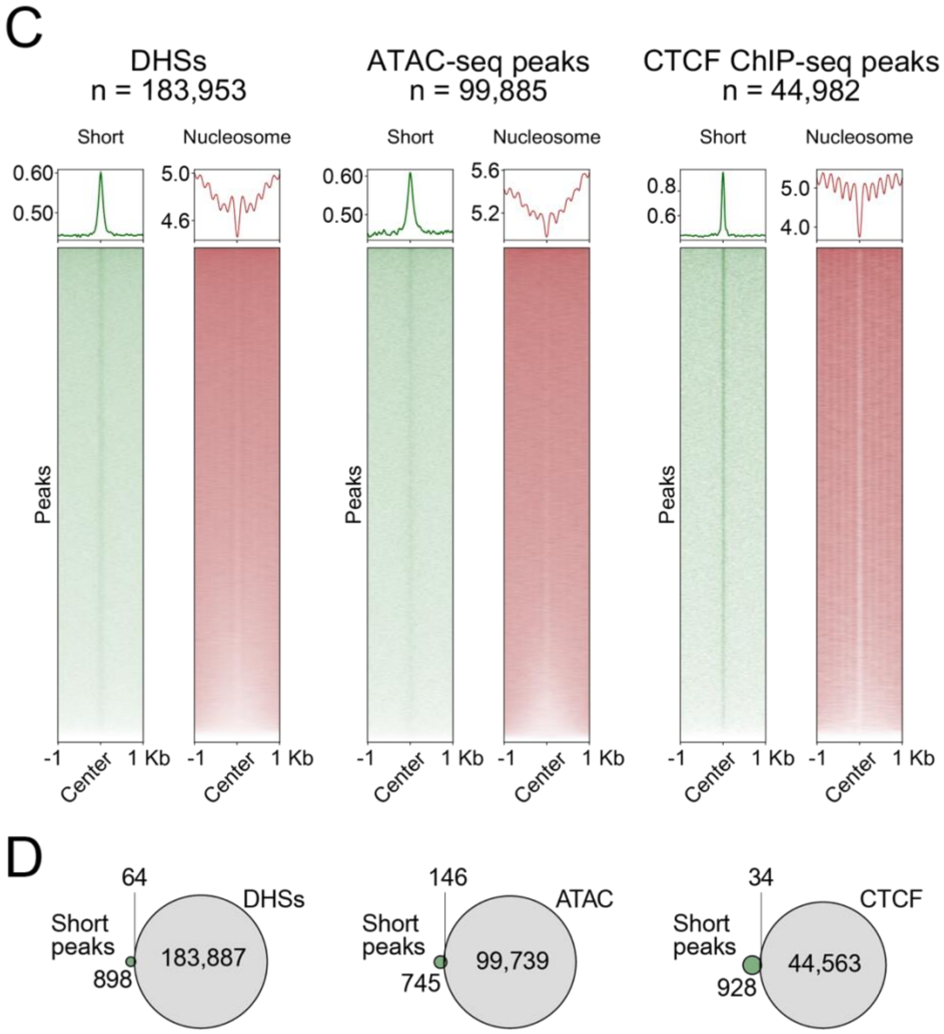


**Supplementary Figure S8. The short single-stranded cfDNA detected by other studies is enriched in the open chromatin regions and TFBS.**

A and C. Aggregation plots of the short (35–80 nt) and nucleosome size range (120–180 nt) single-stranded cfDNA reads presented by Snyder et al. (Snyder et al. 2016) (A) and Burnham et al. (Burnham et al. 2016) (C) on DNaseI-hypersensitive sites (DHSs), the peaks of ATAC-Seq (ATAC), and CTCF binding sites (CTCF).

B and D. The overlapping of the peaks of single-stranded cfDNA ranging from 35 to 80 nt (cfDNA peaks) determined by Snyder et al. (Snyder et al. 2016) (B) and Burnham et al. (Burnham et al. 2016) (D) with DHSs, the peaks of ATAC-Seq, and CTCF binding sites.

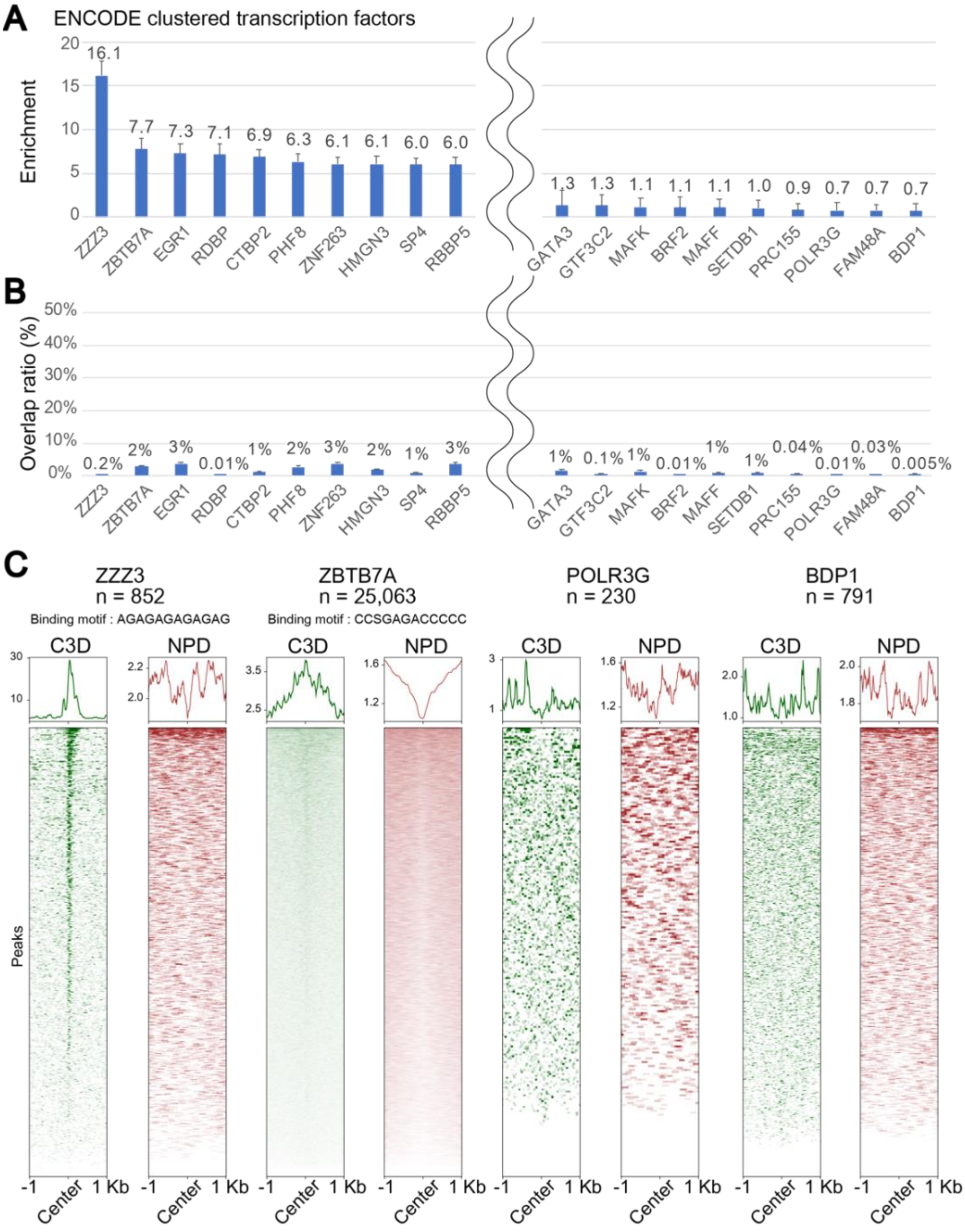


*(Continues on the next page)*

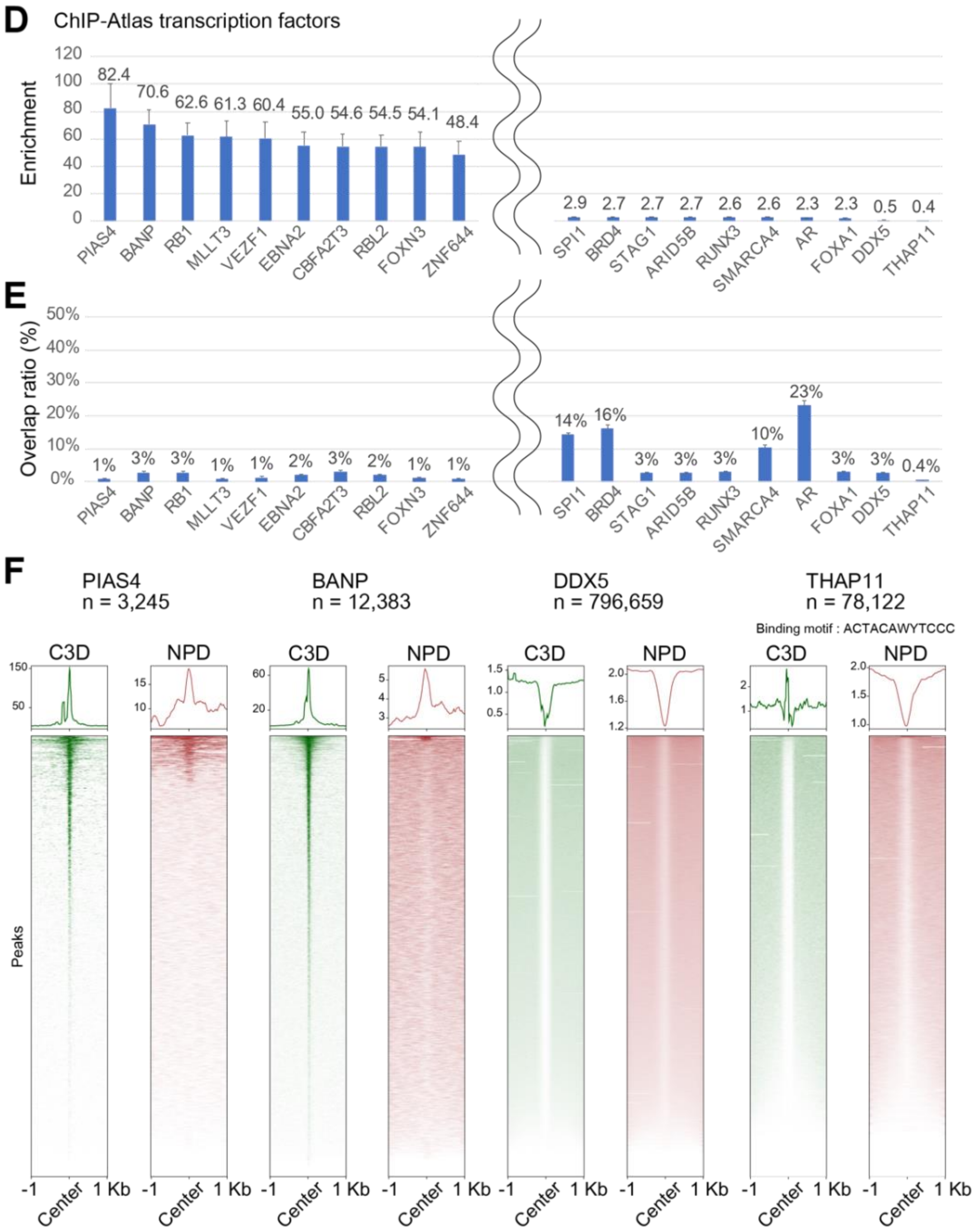


**Supplementary Figure S9. The colocalization of C3D peaks with TFBS.**

The colocalizations of C3D peaks with TFBSs defined by the ENCODE project (A–C) and ChIP-Atlas (D–F) are shown.

A and D. The relative enrichment of C3D peaks on the binding sites of individual TFs over the entire genome was calculated.

B and E. The fractions of the C3D peaks overlapped with those of the individual TFBS.

C and F. Examples of aggregation plots of C3D and NPD reads around the binding sites of individual TFs are shown. Consensus-binding motifs were obtained from ChIPBase (Zhou et al. 2017).

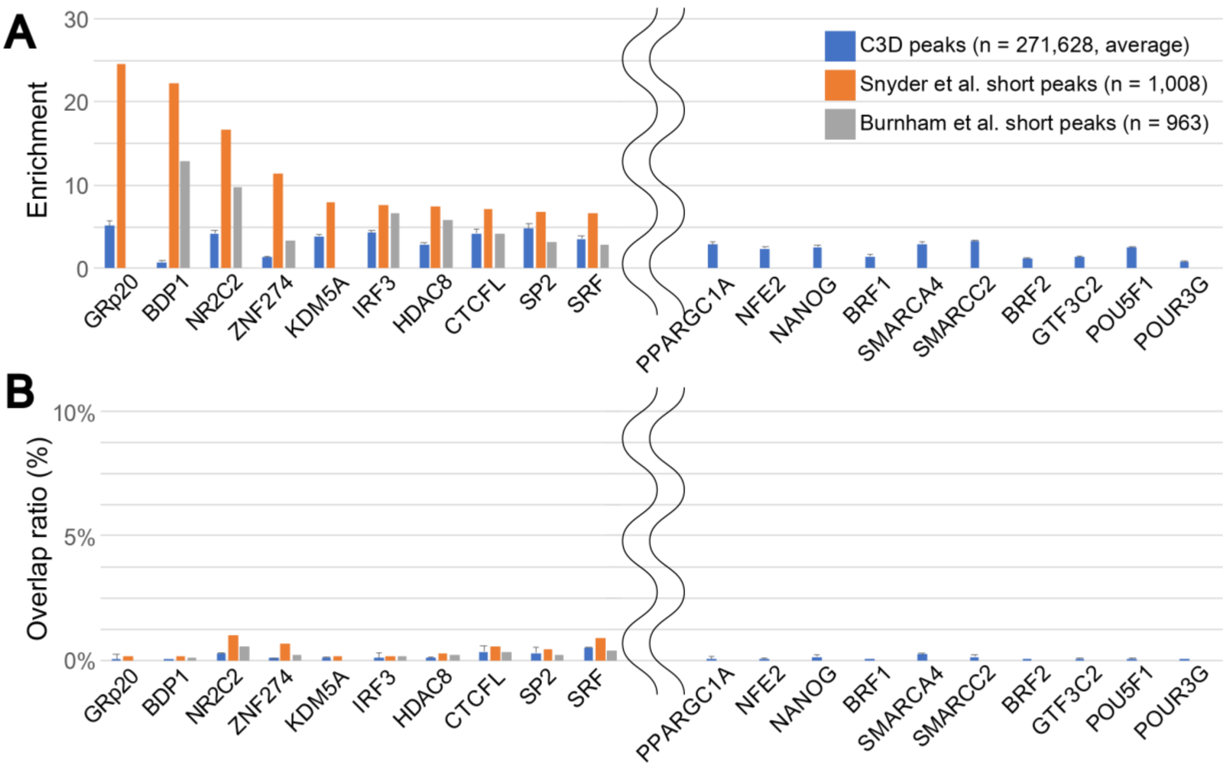


**Supplementary Figure S10. The colocalization of short single-stranded cfDNA with TFBS based on the literature.**

The colocalization of C3D peaks (same as S9), the peaks of the short single-stranded cfDNA (35–80 nt) identified by Snyder et al. (Snyder et al. 2016) and Burnham et al. (Burnham et al. 2016) with TFBSs defined by the ENCODE project are shown.

A. The relative enrichment of the peaks on the binding sites of individual TFs over the entire genome was calculated.

B. The fractions of peaks that overlapped with those of individual TFBS.

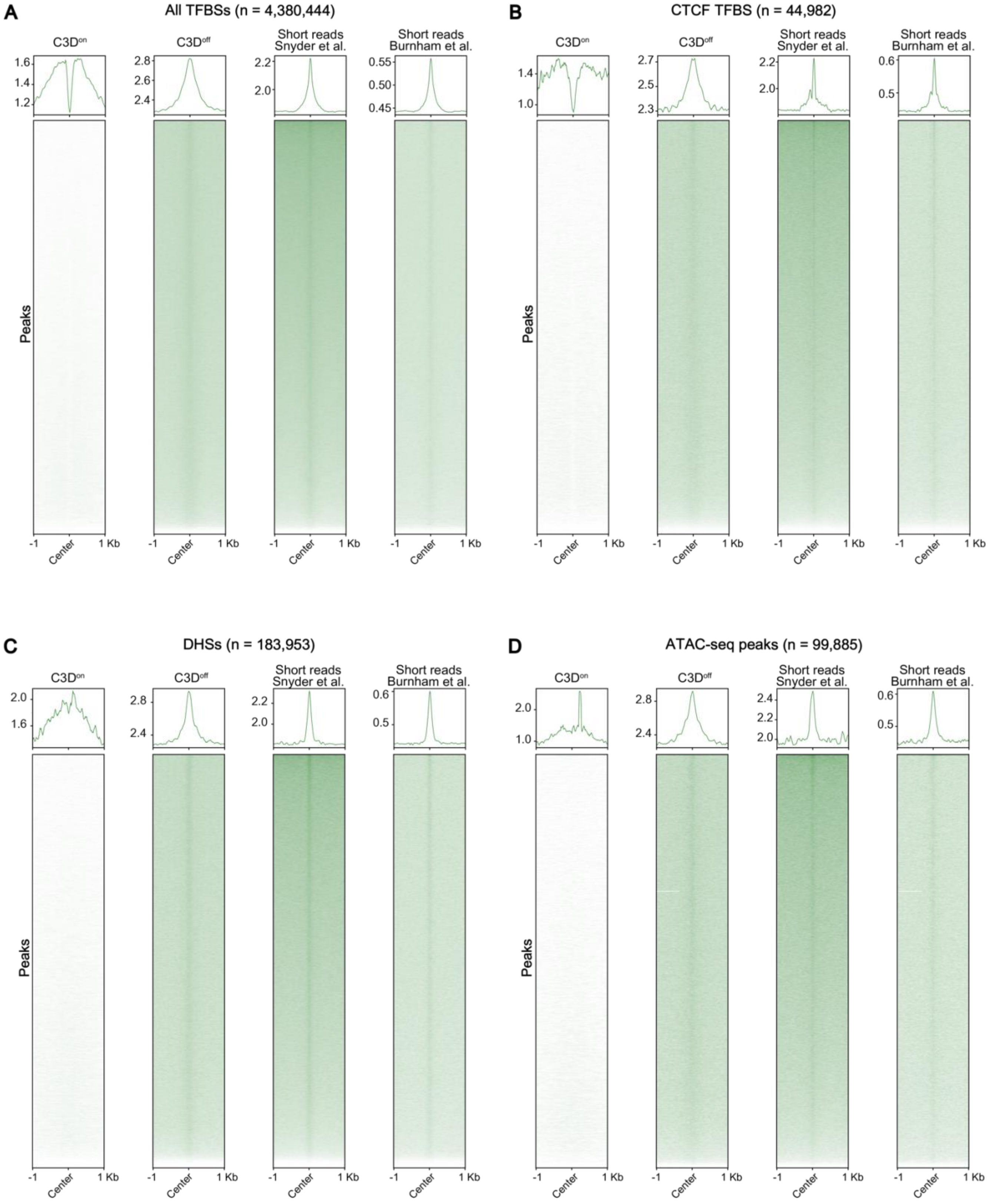


(Continues on the next page)

**Supplementary Figure S11. The colocalization of C3D$^{on}$ and C3D$^{off}$ reads and short single-stranded cfDNA with the open chromatin regions and TFBS based on the literature.** Aggregation plots of C3D$^{on}$ and C3D$^{off}$ reads and short (35–80 nt) cfDNA identified in Snyder et al. (Snyder et al. 2016) and Burnham et al. (Burnham et al. 2016) on all of the transcription factor binding sites (A), CTCF binding sites (B), DHSs (C), and the peaks of ATAC-seq (D) are shown. For all the plots, the TFBS peaks were sorted according to the read number of short cfDNA fragments in Snyder et al. (Snyder et al. 2016).

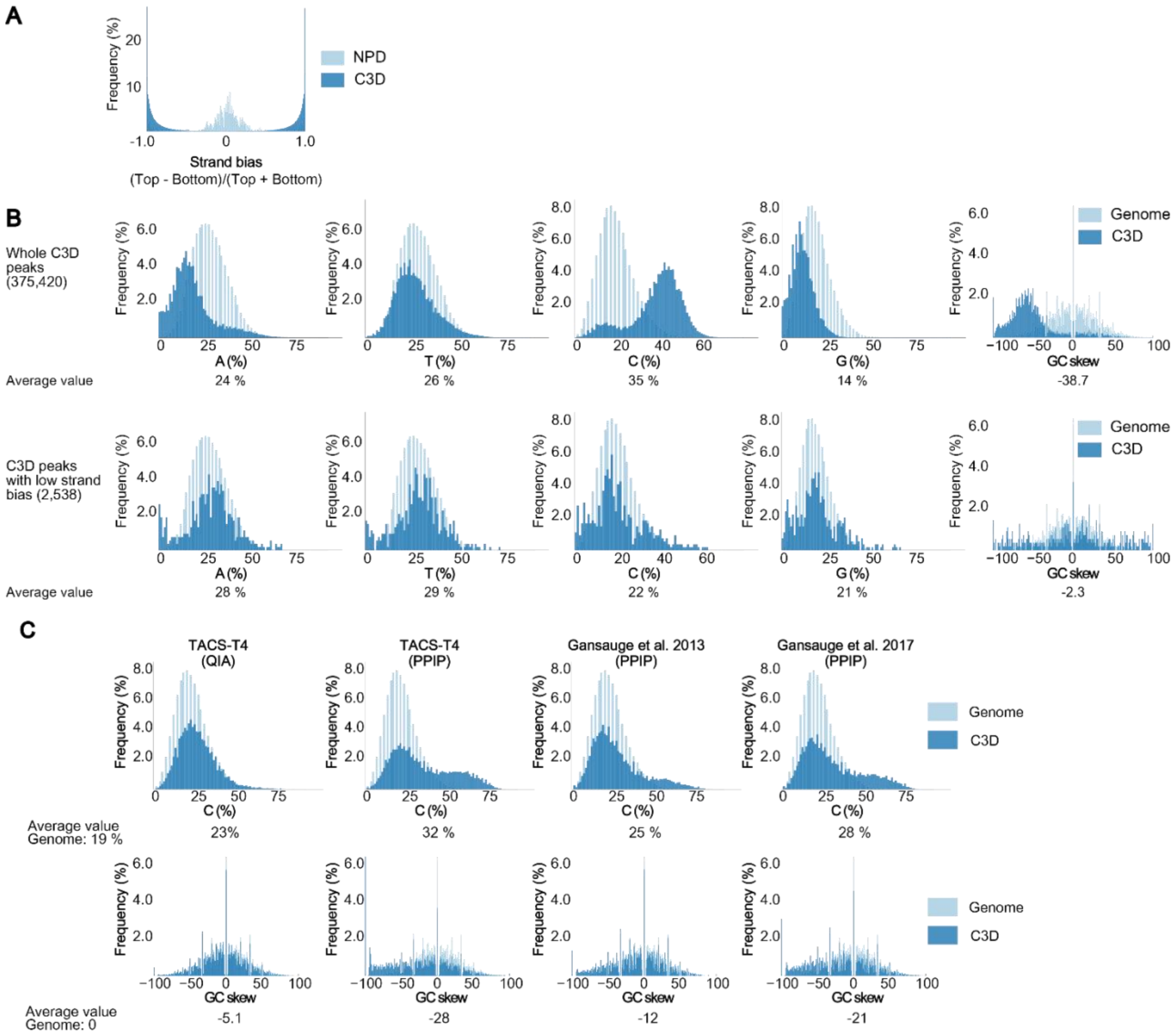


**Supplementary Figure S12. The base composition of C3D peaks was highly biased.**

A. C3D reads are mapped with strong strand biases. The alignments of C3D and NPD reads mapped on the top and bottom strands of the reference genome were served for peak calling by MACS2 separately. Then, the normalized read coverages of the top and bottom strands were calculated for each of the peaks. The strand biases were calculated with the formula at the bottom of the histogram.

B. Histograms display the base composition of C3D peaks (blue) and the randomly chosen 50-mers from the reference genome (light blue). The peaks identified by MACS2 with the reads mapped on the top and bottom strands of the reference genome separately were used for the calculation of the base composition. The distributions of all C3D peaks (top) and peaks with low strand bias (bottom) are shown. If 50% of peak coordinates overlapped with a peak on the opposite strand, the peak was assumed to have less strand bias.

C. Histograms of the C-content and GC-skew for the reads (blue) and randomly chosen 50-mers from the reference genome (light blue). Three different library preparation methods including TACS-T4, Gansuage et al. (2013) (Gansauge and Meyer 2013), and Gausuage et al. (2017) (Gansauge et al. 2017) are compared for the short reads (35–75 bp). As a control, a library prepared with TACS-T4 library from cfDNA purified with the QIAamp Circulating Nucleic Acid Kit (Qiagen) was also calculated.

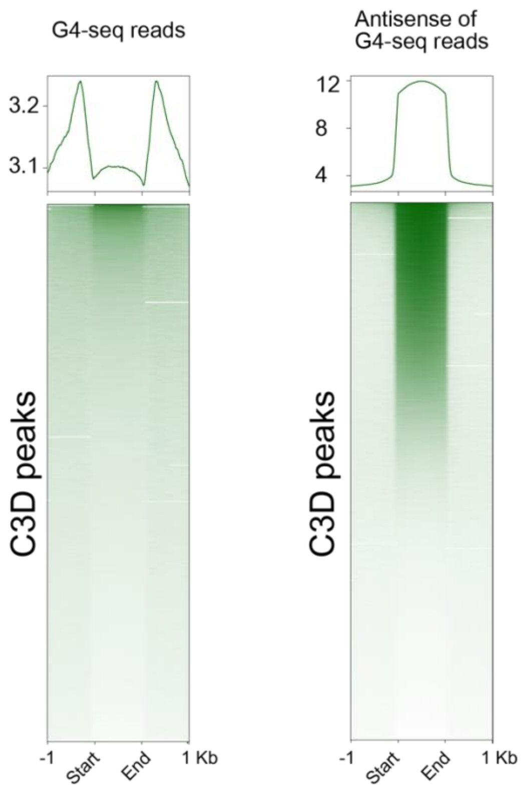


**Supplementary Figure S13. The antisense strand of G4-Seq reads well localize on the C3D peaks.**

Aggregation plots of G4-seq read coverage of the C3D peaks. The normalized read coverage of G4-seq (left) and that of the opposite strand (right) are shown.

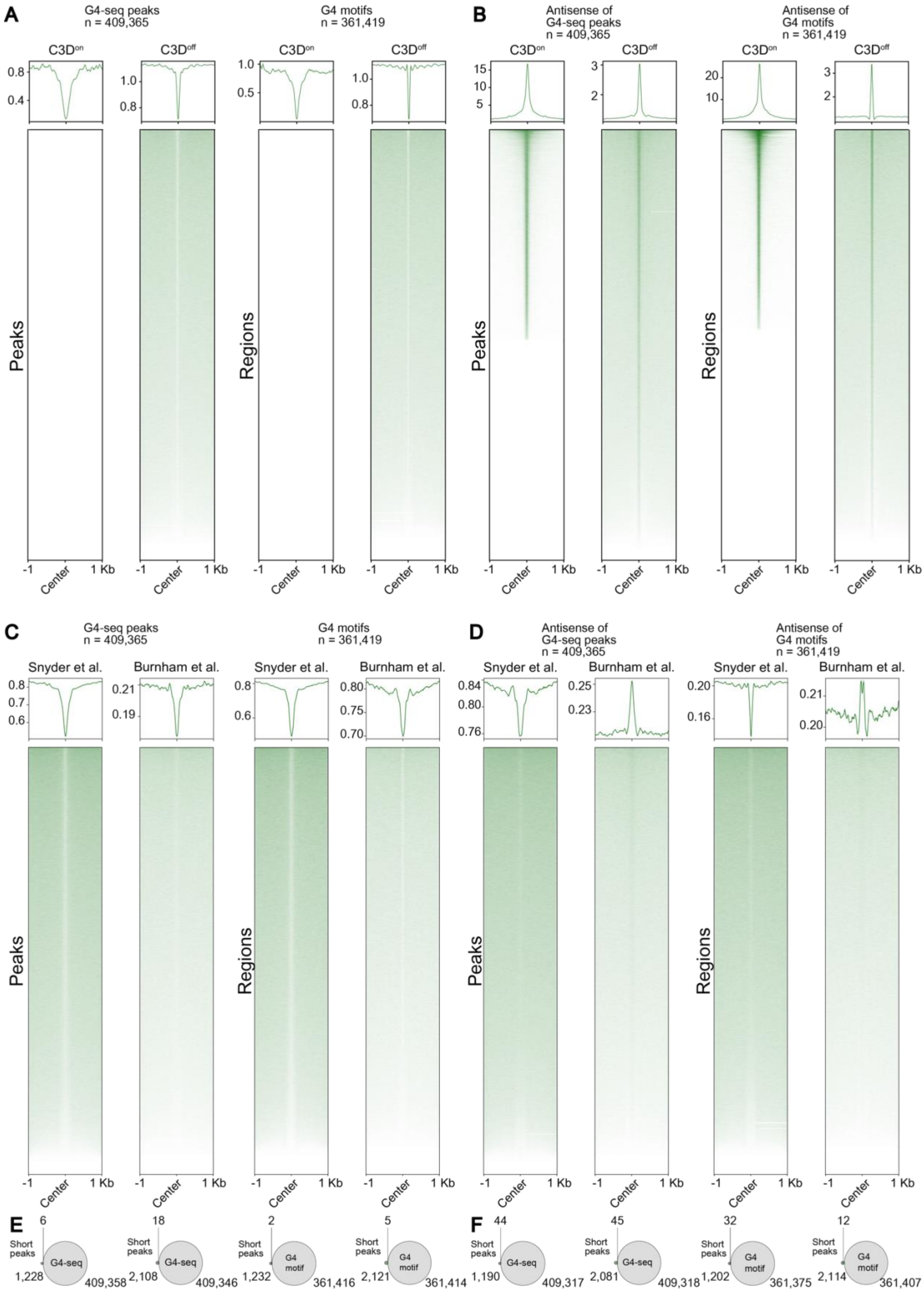


(Continues on the next page)

**Supplementary Figure S14. Colocalization analysis of the G4 structure with short single-stranded cfDNA in the literature as well as C3D$^{on}$ and C3D$^{off}$ reads.**

A and B. Aggregation plots of C3D$^{on}$ and C3D$^{off}$ reads on the same (A) and antisense (B) strands of G4-seq peaks (left) and putative G4 sequences defined by the *quadparser* algorithm (right).

C and D. Aggregation plots of short (35–80 nt) single-stranded cfDNA reads presented by Snyder et al. (Snyder et al. 2016) and Burnham et al. (Burnham et al. 2016) on the same (C) and antisense (D) strands of G4-seq peaks (left) and putative G4 sequences defined by the *quadparser* algorithm (right).

E and F. Venn diagrams showing the overlapping of the peaks of the short (35–80 nt) single-stranded cfDNA presented by Snyder et al. (Snyder et al. 2016) and Burnham et al. (Burnham et al. 2016) on the same (E) and antisense (F) strands of G4-seq peaks (left) and putative G4 sequences identified by the *quadparser* algorithm (right).

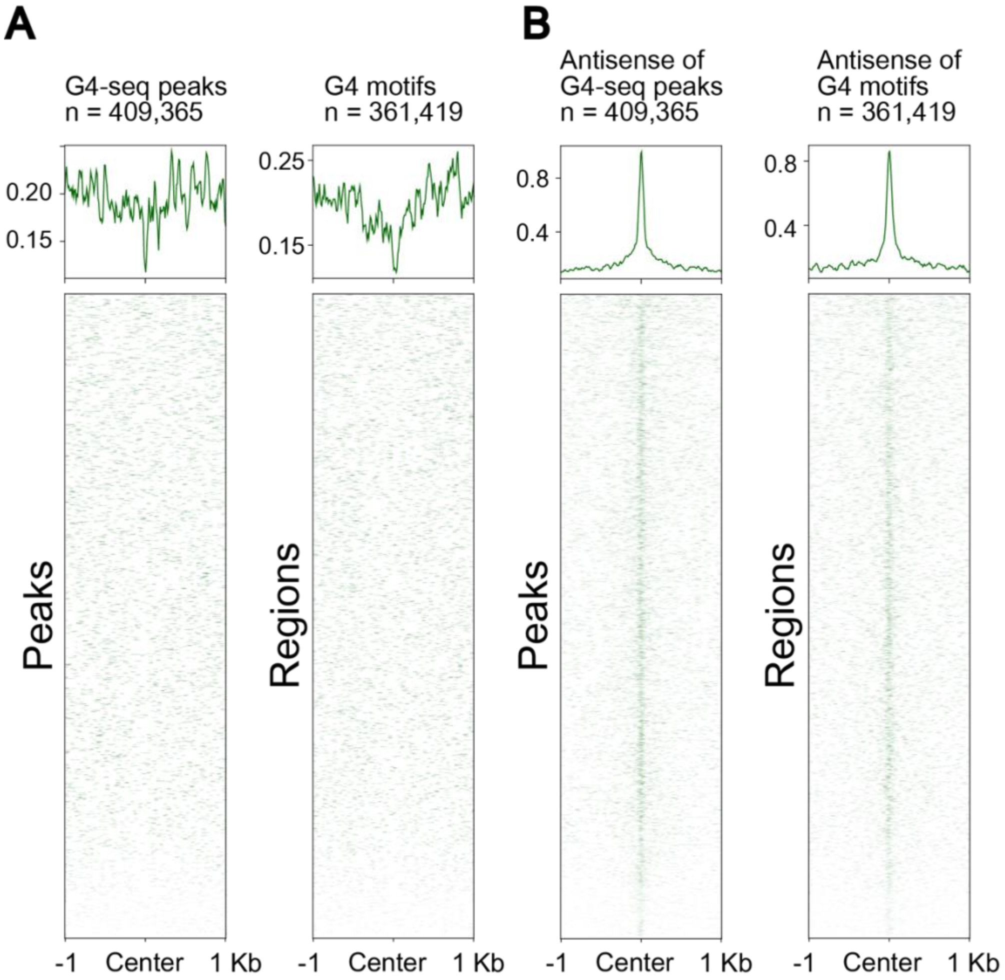


*(Continues on the next page)*

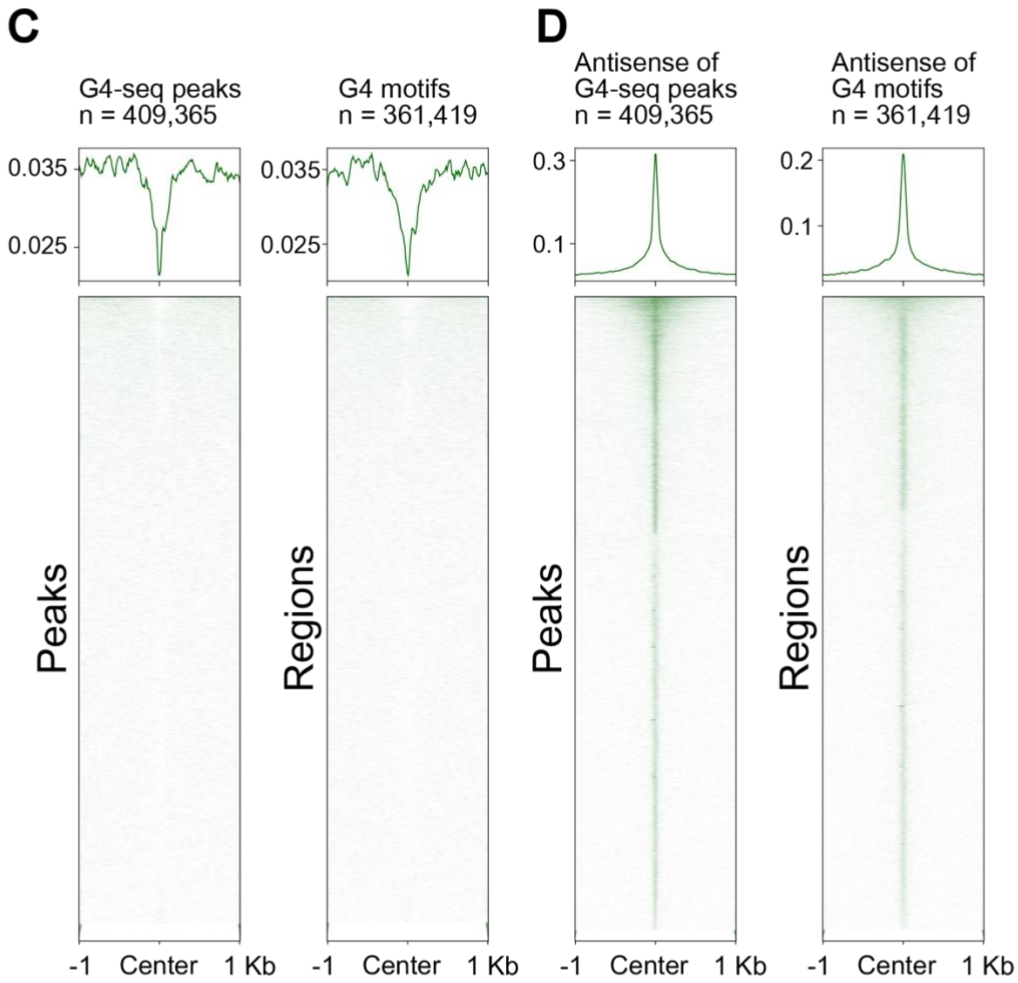


**Supplementary Figure S15. Enriched C3D reads on the G4 structure with different library preparation methods.**

The aggregation plots for the C3D reads obtained with the libraries prepared by Gansauge et al. (2013) (Gansauge and Meyer 2013) (A and B) and Gansauge et al. (2017) (Gansauge et al. 2017) (C and D) were drawn using the same (A and C) and antisense (B and D) strands of G4-seq peaks and putative G4 sequences defined by the *quadparser* algorithm.

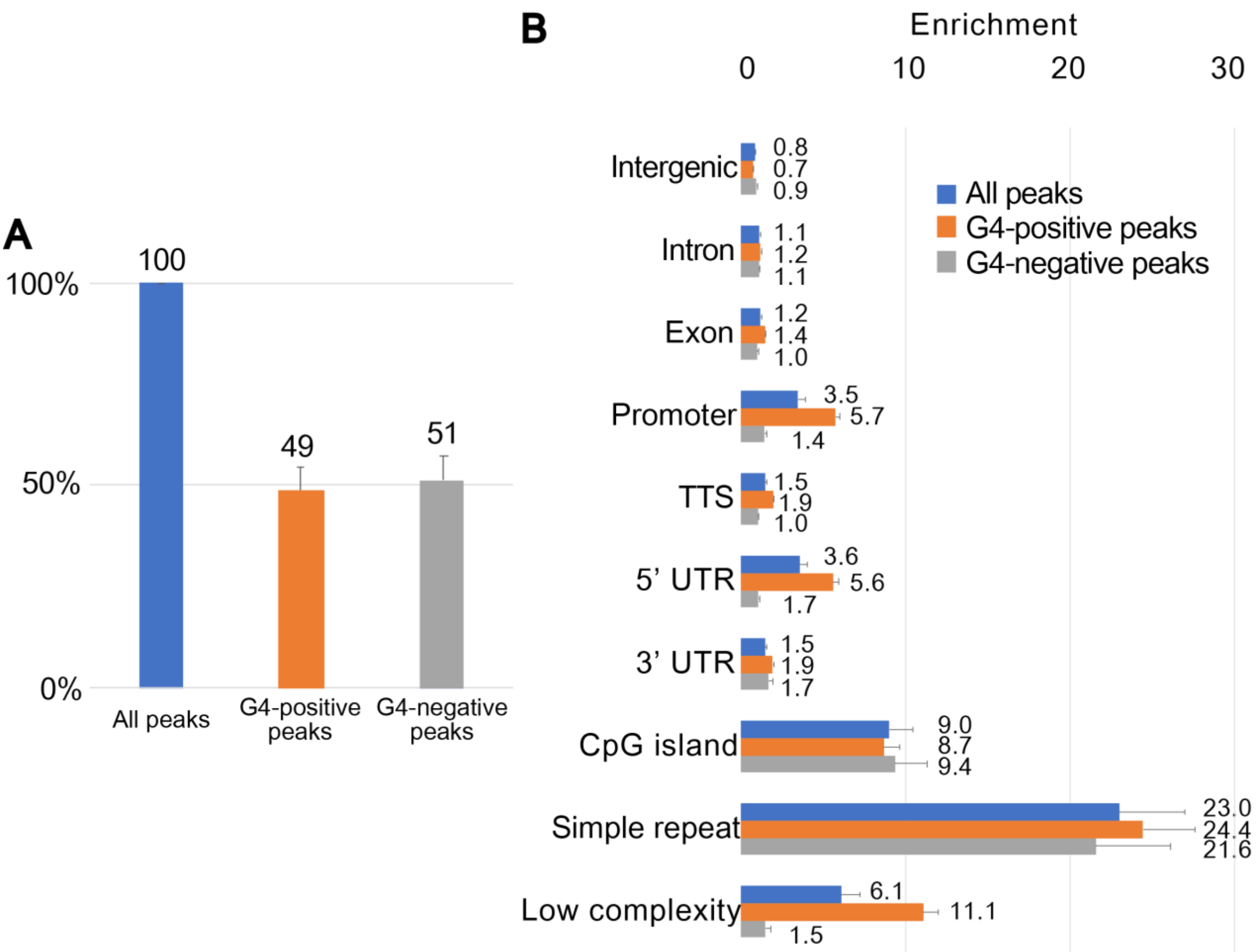


**Supplementary Figure S16. C3D peaks without G4 regions are less enriched in the regulatory regions.**

A. The fractions of G4-positive C3D peaks (C3D peaks overlapping with either G4-seq peaks or G4 motifs) and G4-negative peaks (C3D peaks without overlap from either G4-seq peaks or G4 motifs). The average values of the five donors are shown.

B. The enrichment analysis for all of the C3D peaks (same as Figure 5B), G4-positive peaks, and G4-negative peaks were similarly performed as described for Figure 5B.

**Supplementary Tables**

**Supplementary Table S1. Summary of the library preparations and sequencing methods**

| Sample ID | DNA purification method[*1] | Library preparation | | | | | Sequencing Platform | Number of reads[*3] | | | | Number of peaks | | Percentages of reads in peaks[*4] | |
|---|---|---|---|---|---|---|---|---|---|---|---|---|---|---|---|
| | | Pretreatment[*2] | Protocol | Input DNA (ng) | PCR cycles | Yield (pmol) | | Total | Uniquely mapped | C3D | NPD | C3D | NPD | C3D | NPD |
| D1 | PPIP | Heat | TACS-T4 | 6.6 | 5 | 105 | MiSeq | 643K | 448K | 194K | 140K | NA | NA | NA | NA |
| D1 | QIA | Heat | TACS-T4 | 3.9 | 5 | 26 | MiSeq | 351K | 254K | NA | 128K | NA | NA | NA | NA |
| D1 | PPIP | Exo | TACS-T4 | 8.1 | 5 | 64 | MiSeq | 574K | 443K | NA | 158K | NA | NA | NA | NA |
| D1 | PPIP | | ThruPlex | 6.6 | 5 | 126 | MiSeq | 660K | 514K | NA | 330K | NA | NA | NA | NA |
| D1 | PPIP | | TACS-T4 | 6.6 | 5 | 16 | MiSeq | 573K | 356K | 233K | 62K | NA | NA | NA | NA |
| D1 | PPIP | | TACS-T4 | 8.6 | 5 | 16.9 | HiSeq X | 523M | 266M | 143M | 72M | 375K | 493 | 25% | 0.3% |
| D2 | PPIP | | TACS-T4 | 11 | 5 | 9.4 | HiSeq X | 531M | 250M | 176M | 70M | 284K | 398 | 18% | 0.2% |
| D3 | PPIP | | TACS-T4 | 11 | 5 | 2.2 | HiSeq X | 533M | 255M | 205M | 50M | 250K | 342 | 21% | 0.1% |
| D4 | PPIP | | TACS-T4 | 12 | 5 | 3.3 | HiSeq X | 544M | 264M | 184M | 90M | 186K | 337 | 15% | 0.2% |
| D5 | PPIP | | TACS-T4 | 8.6 | 5 | 8.6 | HiSeq X | 536M | 193M | 114M | 83M | 261K | 365 | 25% | 0.2% |
| D1 | PPIP | Heat | TACS-T4 | 21 | 10 | 580 | Miseq | 1.6M | 1.1M | 402K | 414K | NA | NA | NA | NA |
| D1 | PPIP | Heat | Gansauge et al. (2013) (Gansauge and Meyer 2013) | 21 | 16 | 2.2 | Miseq | 0.76M | 0.42M | 223K | 93K | NA | NA | NA | NA |
| D1 | PPIP | Heat | Gansauge et al. (2017) (Gansauge et al. 2017) | 21 | 13 | 88 | Miseq | 1.3M | 0.93M | 525K | 225K | NA | NA | NA | NA |

*1 “PPIP” means that the procedure sequentially performed proteinase K treatment, phenol, and phenol-chloroform extractions, and isopropanol precipitation, as described in the Materials and Methods section. “QIA” means QIAamp Circulating Nucleic Acid Kit (Qiagen).
*2 “Heat” denotes heat denaturation. “Exo” means exonuclease (exonuclease I and RecJf) treatment.
*3 K and M denote thousands and millions, respectively.
*4 Percentages of C3D and NPD reads that overlap with the C3D and NPD peaks, respectively.
NA denotes “Not Applicable”.

**Supplementary Table S2. The number of reads and peaks calculated from two publicly available single-stranded cfDNA datasets**

| Dataset name | DNA purification method*1 | Sequencing | | Number of reads*2 | | | | Number of peaks | | Percentage of reads in peaks*3 | |
|---|---|---|---|---|---|---|---|---|---|---|---|
| | | Library preparation method | Platform | Total | Uniquely mapped | Short*4 | Nucleosome*4 | Short*4 | Nucleosome*4 | Short*4 | Nucleosome*4 |
| Snyder et al. (Snyder et al. 2016) | QIA | Gansauge et al. (2013) (Gansauge and Meyer 2013) | HiSeq 2000 and NextSeq 500 | 638M | 473M | 80M | 249M | 1,008 | 774 | 0.5% | 0.3% |
| Burnham et al. (Burnham et al. 2016) | QIA | Gansauge et al. (2013) (Gansauge and Meyer 2013) | MiSeq and HiSeq | 207M | 154M | 19M | 94M | 963 | 379 | 0.4% | 0.3% |
| This study (Mean value) | PPIP | TACS-T4 | HiSeq X | 533M | 246M | 165M | 73M | 271,628 | 387 | 21% | 0.2% |

*1 "PPIP" means the procedure sequentially performed proteinase K treatment, phenol, phenol-chloroform extractions, and isopropanol precipitation, as described in the Materials and Methods section. "QIA" means QIAamp Circulating Nucleic Acid Kit (Qiagen).
*2 M denotes million.
*3 Percentage of reads overlapping with the peaks.
*4 "Short" means 35–80 nt reads. "Nucleosome" indicates the nucleosome size range (120–180 nt) reads from previous studies. The mean values of C3D and NPD reads are shown in this study.

**Supplementary Tables S3, S4, and S5 have been supplied as Excel spreadsheets.**

**Supplementary Table S6. Blood samples used in the current study**

| Sample ID | Sample type | Age (years), Sex | Supplier[*1] |
|---|---|---|---|
| D1 | Healthy donor | 32, Male | |
| D2 | Healthy donor | 30, Male | |
| D3 | Healthy donor | 43, Male | |
| D4 | Healthy donor | 44, Male | |
| D5 | Healthy donor | 35, Male | |
| P1 | Purchased plasma | 21, Female | BIOPREDIC |
| P2 | Purchased plasma | 48, Male | BIOPREDIC |
| P3 | Purchased plasma | 52, Male | BIOPREDIC |
| S1 | Purchased serum | 61, Female | CTLS |
| S2 | Purchased serum | 29, Male | Cosmo bio |

*1 BIOPREDIC, CTLS and Cosmo bio indicate that we obtained these samples from BIOPREDIC (Rennes, France), Cosmo Bio (Tokyo, Japan), and Clinical Trials Laboratory Services (London, UK).

**Supplementary Table S7. Oligonucleotides used in the current study**

| Name | Used for | Nucleotide sequence and chemical modifications |
|---|---|---|
| PA-Truseq Index-dSp-P | TACS ligation | 5'-[phosphate] AGATCGGAAGAGCACACGTCTGAACTCCAG [dSpacer] CAC [phosphate]-3' |
| TruseqUniv-T | Complementary strand synthesis | 5'-TCTTTCCCTACACGACGCTCTTCCGATCTT-3' |
| PCR-Univ | Indexing and amplification | 5'-AATGATACGGCGACCACCGAGATCTACACTCTTTCCCTACACGAC-3' |
| PCR-Index-X[*1] | Indexing and amplification | 5'-CAAGCAGAAGACGGCATACGAGAT [Index Sequence] GTGACTGGAGTTCAGACGT-3' |
| N [Number][*2] | Size marker and model substrate | 5'-N[Number]-3' |
| rA40[*3] | Control for RNase digestion | 5'-[rA]40-3' |
| 200 bp-forward | PCR primer for dsDNA control | 5'-CGACCTTTCTGTGGTGAA-3' |
| 200 bp-reverse | PCR primer for dsDNA control | 5'-CACATGTCGCGGTGGTTA-3' |

*1 “X” is the index number, i.e., a specific hexamer was inserted in the position “[Index Sequence]” for indexing sequences (see Supplementary Table S8).

*2 N denotes an equimolar mixture of A, C, G, and T. “[Number]” refers to the nucleotide length. In the current study, stretches of 40, 60, 80, 100, 120, 140, and 160 nt were used.

*3 [rA] denotes an adenosine residue.

**Supplementary Table S8. Indexing sequences**

| Index number[*1] | Index sequences[*2] |
|---|---|
| 1 | 5'-CGTGAT-3' |
| 2 | 5'-ACATCG-3' |
| 3 | 5'-ACATCG-3' |
| 4 | 5'-TGGTCA-3' |
| 5 | 5'-CACTGT-3' |

*1 This is the same as “X” used in Supplementary Table S7.
*2 This is the “[Index Sequence]” used in Supplementary Table S7.

**Supplementary Table S9. Sources of publicly available data used in the current study**

| Data name | Reference | Accession number (Database) |
|---|---|---|
| FANTOM5 enhancers and promoters | Andersson et al. (Andersson et al. 2014) | |
| RefSeq genes | O'Leary et al. (O'Leary et al. 2016) | - |
| ATAC-seq of GM12878 | Buenrostro et al. (Buenrostro et al. 2013) | GSE47753 (NCBI GEO) |
| ENCODE TFBS cluster version 3 | Gerstein et al. (Gerstein et al. 2012) | - |
| ChIP-Atlas | Oki et al. (Oki et al. 2018) | - |
| G4-seq reads and peaks | Chambers et al. (Chambers et al. 2015) | GSE63874 (NCBI GEO) |
| CpGi from UCSC table browser | Karolchik et al. (Karolchik et al. 2004) | - |
| Snyder et al. (Snyder et al. 2016) | Snyder et al. (Snyder et al. 2016) | GSE71378 (NCBI GEO) |
| Burnham et al. (Burnham et al. 2016) | Burnham et al. (Burnham et al. 2016) | PJRNA306662 (NCBI SRA) |

**Supplementary Table S10. Files of the ENCODE datasets downloaded from the UCSC Table Browser**

| Experiment type | Cell line | BigWig file | BED file |
|---|---|---|---|
| DNase-seq | GM12878 | wgEncodeOpenChromDnaseGm12878Sig.bigWig | wgEncodeOpenChromDnaseGm12878Pk.narrowPeak |
| CTCF ChIP-seq | GM12878 | wgEncodeUwTfbsGm12878CtcfStdRawRep1.bigWig | wgEncodeUwTfbsGm12878CtcfStdPkRep1.narrowPeak.gz |
| MNase-seq | GM12878 | wgEncodeSydhNsomeGm12878Sig.bigWig | - |

### Supplementary Methods

**Preparation of model dsDNA fragments**

We prepared a model double-stranded DNA fragment (DNA200). A 50-µL reaction containing 1× PrimeStar Max (Takara Bio Inc.), 0.2 µM each of 200 bp-forward and 200 bp-reverse (Supplementary Table S7), and 5 ng of lambda DNA was prepared. Then, PCR amplification was performed with the following conditions: initial denaturation at 95°C for 1 min; 35 cycles of 2-step incubation at 95°C for 15 sec and 72°C for 30 sec; final extension at 72°C for 5 min. Following the manufacturer's instructions, the amplified DNA was purified with AMPure XP (Beckman Coulter, Brea, CA, USA).

**Figure 1C and Supplementary Figure S1D**

A model mixture was prepared by mixing equimolar amounts of the synthetic DNA, N40, N80, N120, and DNA200 at 0.1 µM in 10 mM Tris-HCl (pH 8.0). Next, 5 µL of purified cfDNA, which was equivalent to 0.25 mL of plasma, was supplemented with 0.2 µL of the control mixture, and subjected to 3′-terminal fluorescent labeling and analyzed as described in Materials and Methods.

**Figure 1D and Supplementary Figure S1E**

A model mixture was prepared by mixing equimolar amounts of the synthetic DNA N60, N80, N100, N120, N140, N160, synthetic RNA rA40 (see Supplementary Table S7), and DNA200 at 0.05 µM in 10 mM Tris-HCl (pH 8.0). In 2 µL of the mixture or 10 µL of cfDNA extracted from 1 mL plasma, 1.2 µL 10× TACS buffer (500 mM HEPES-KOH (pH 7.5), 50 mM $MgCl_2$, and 5% (v/v) Triton-X100), and 1.2 µL 10× phosphate buffered saline (PBS) were supplemented, and the reaction volume was adjusted to 12 µL. Nuclease digestion was started with the addition of 15 units of RNase If (New England Biolabs; NEB, Ipswich, MA, USA), 1 unit of RQ1 RNase-Free DNase (Promega, Fitchburg, WI, USA), or a mixture containing both six units of exonuclease I and ten units of RecJf (NEB). The reaction mixture was incubated at 37°C for 30 min, diluted in 1 mL PBS, and purified again using the PPIP method. The purified DNA was analyzed as described in the Materials and Methods.

**Figure 1E and Supplementary Figure S1F**

First, a tube containing 0.5 mL of plasma was supplemented with 5 µL of 1 M $MgCl_2$. If indicated, 1 µL of the model mixture, as used in Figure 1D, was spiked into the plasma. Then, 50 units of RNase If (NEB), 20 units of RQ1 RNase-Free DNase (Promega), or a mixture containing 20 units of exonuclease I and 30 units of RecJf were added. The reaction was incubated at 37°C for 30 min and was then purified using the PPIP method. The purified cfDNA was analyzed via denaturing gel electrophoresis, as described in Materials and Methods.

**Supplementary Figure S1B, S1C, and S2B**

A control mixture was prepared by mixing N40, N80, N120, and DNA200 at 0.1 µM in 10 mM Tris-HCl (pH 8.0). To 500 µL of plasma or serum, 0.4 µL of the control mixture was added, and the cfDNA was purified, as described in Materials and Methods. The purified DNA was analyzed as described in the Materials and Methods. The amounts of the supernatants or pellets after ultracentrifugation equivalent to 500 µL of plasma were used for the results presented in Supplementary Figure S2B.

**Figure 2E and Supplementary Figure S4D**

Plasma (0.5 mL), 1 M $MgCl_2$ (5 µL), exonuclease I (1 µL), and RefJf (1 µL) were combined and incubated at 37°C for 30 min. Then, cfDNA was purified as described in the Materials and Methods section. The purified cfDNA was then used for the TACS-T4 scheme, as described in Materials and Methods.

**Figure 2G and Supplementary Figure S4F**

For the library "without heat denaturation," TACS ligation was performed without any pretreatment of cfDNA, as follows. A reaction was prepared by mixing 10 µL of cfDNA extracted from 0.5 mL plasma, 2.5 µL of 10× TACS Buffer, 10 µL of 50% (w/v) PEG, 1 µL of 10 µM PA-TruSeqIndex-dSp-P (Supplementary Table S7), 1 µL of 10 mM ATP, 1 µL of TdT, and 1 µL of 2 mg/mL TS 2126 RNA ligase in a 25 µL volume and sequentially incubated at 37°C for 30 min, 65°C for 2 h, and 95°C for 5 min. After this step, library preparation was performed as described in the Materials and Methods section.

**Expression and purification of TS2126 RNA ligase**

The previously used plasmid encoding TS2126 RNA ligase (19; Addgene #119941) was modified with inverse PCR so that the recombinant protein had a Strep-Tag at its carboxyl-terminal (Addgene #159350). The plasmid was then used to transform T7 Express *E. coli* (NEB). Transformants were inoculated into 3 mL of 2× YT medium (1.6% [w/v] Bacto Tryptone, 1% [w/v] Bacto yeast extract, 0.5% glucose, and 0.5% [w/v] NaCl) supplemented with 50 µg/mL kanamycin. They were grown at 37°C with continuous shaking at 250 rpm for

16–18 h. The cultures were diluted in 1 L of 2× YT medium containing 50 µg/mL of kanamycin and incubated at 37°C with continuous shaking at 120 rpm for 4–5 h. Protein expression was induced by the addition of isopropyl-β-D-thiogalactopyranoside (IPTG) powder (Nacalai Tesque, Kyoto, Japan) at a final concentration of 1 mM, followed by further incubation at 37°C with continuous shaking at 120 rpm for 4 h. Cells were collected via centrifugation, suspended in 20 mL of HisTrap Buffer A (50 mM sodium phosphate, pH 7.0, 100 mM NaCl, and 10 mM imidazole), and then lysed via sonication using a Digital Sonifier (Branson, Danbury, CT, USA). The cell debris was removed by centrifugation at 10,000 × *g* for 10 min, followed by filtration using 32-mm Acrodisc syringe filters with Supor Membrane (0.45-µm) (Pall, Port Washington, NY, USA). The cleared cell lysate was loaded onto the AKTA start chromatography system (GE Healthcare, Chicago, IL, USA) equipped with a HisTrap HP column (5-mL, GE Healthcare) using an equipped template for affinity chromatography. The column was equilibrated and washed with HisTrap Buffer A, and the purified protein was eluted with HisTrap Buffer B (50 mM sodium phosphate, pH 7.0, 100 mM NaCl, and 200 mM imidazole). Fractions containing the target protein were combined, diluted 20-fold with Strep buffer A (100 mM Tris-HCl, pH 8.0, 100 mM NaCl, and 1 mM EDTA), and further purified using the AKTA start system equipped with a StrepTrap HP column (5 mL, GE Healthcare). StrepTrap purification was performed using an equipped template for affinity chromatography with Strep Buffer A and Strep Buffer B (100 mM Tris-HCl, pH 8.0, 100 mM NaCl, 1 mM EDTA, and 2.5 mM d-Desthiobiotin). Fractions containing the target protein were combined and subjected to ultra-filtration on an Amicon-4 30K device (Millipore, Burlington, MA, USA) to exchange the buffer with 2× storage buffer (100 mM Tris-HCl [pH 8.0], 150 mM NaCl, 1 mM EDTA) and concentrate the protein. Protein concentration was determined using the Protein Assay CBB Solution (Nacalai Tesque). After adjusting the protein concentration to 4 mg/mL, the protein solution was combined with an equal volume of glycerol and stored at -20°C until use.

**Expression and purification of recombinant human aprataxin**

DNA fragments encoding human aprataxin (UniProt#Q7Z2E3) were chemically synthesized by Eurofins Genomics (Tokyo, Japan) with codon optimization for *Escherichia coli*. Recognition sequences for BamHI and EcoRI were introduced at the 5' and 3' ends of the protein-coding sequence, respectively. The fragment was subcloned into the BamHI–EcoRI site of pColdI (Takara Bio Inc.). The obtained plasmid was then used to transform *E. coli* BL21 (Takara Bio Inc.). The transformants were inoculated into 3 mL of 2× YT medium containing

50 μg/mL of carbenicillin. They were grown at 37°C with continuous shaking at 250 rpm for 16–18 h. Next, the cultures were diluted in 1 L of 2× YT medium containing 50 μg/mL of carbenicillin and grown with shaking at 37°C for 4–5 h. The culture was then cooled to 16°C in ice-cold water for 30 min. Protein expression was induced by the addition of IPTG powder to obtain a final concentration of 1 mM. The culture was then incubated with shaking at 16°C for at least 16 h. The first purification using the HisTrap HP column was performed as described for the purification of TS2126 RNA ligase. Fractions containing the target protein were combined, diluted 20-fold with heparin buffer A (50 mM Tris-HCl, pH 8.0), and further purified using the AKTA start system equipped with a HiTrap Heparin HP column (GE Healthcare). The second purification was performed using an equipped template for ion-exchange chromatography with heparin buffers A and B (50 mM Tris-HCl [pH 8.0], and 1 M NaCl). The subsequent buffer exchange was performed using the Amicon-4 30K device, as described for the purification of TS2126 RNA ligase. After adjusting the protein concentration to 2 mg/mL, the solution was combined with an equal volume of glycerol and stored at -20°C until use.

**Figure 3**

First, bowtie2 was used to map the sequenced reads on the human reference genome, including those for nuclear and mitochondrial sequences. The unmappable reads of the human genome were compared with the bacterial genome by using NCBI BLAST (Altschul et al. 1990) and reference databases comprising viral, bacterial, and fungal genome sequences (downloaded from NCBI ftp://ftp.ncbi.nih.gov/genomes on April 17, 2019), as described by Vlaminck et al. (De Vlaminck et al. 2013).

**Figure 4C**

The normalized read coverage of C3D peaks for the five donors (Supplementary Table S1) was converted to a matrix with deepTools multiBigwigSummary and plotted with plotCorrelation (Ramirez et al. 2016).

**Supplementary Figure S7B**

GO Enrichment analysis was performed by PANTHER (Thomas et al. 2003), and GOslim was used as a GO term set. Fisher's exact test was performed against all reference genes using R.

**Pipelines for bioinformatic analyses**

The commands used in the bioinformatic analyses are shown in Supplementary Information S3, with flowcharts of analytical pipelines.

### Supplementary Information

**Supplementary Information S1. Nucleotide sequence of a gene encoding codon-optimized TS2126 RNA ligase with a Strep-Tag.**

Recognition sequences for BamHI and EcoRI are underlined.
The sequence encoding the Strep-Tag is shown in bold.

GGATCCATGAGCTCACTGGCTCCGTGGCGTACGACGAGCTGGAGTCCGCTGGGCTCTCCGCC
AAGTTTAGAGGATGCTTTGCGTCTTGCGCGCACAACTCGCGCATTCGCAGTCCGCCGCGATG
GTGAAGGTCGCGCATTGGTTACCTACCTGTATGGCACTCCCGAGCTGTTCTCCCTGCCGGGC
GCGCGTGAATTGCGTGGTATCGTGTATCGCGAGGAGGATGGCACCGTGCTGAGCCGTCCGTT
TCACAAATTCTTCAACTTTGGAGAACCGTTAGCTCCGGGTGAAGAGGCCTTTAAAGCATTTC
GCGATTCGATGGTGCCCCTGTTTGTCGCCGAGAAAGTGGATGGCTACCTGGCACAAGCGTAC
CTGGATGGTGGGGAGTTGCGTTTTGCCTCTCGGCATAGCCTTAATCCGCCACTTGTGGGTGC
GTTGCTGCGCAAAGCCGTCGATGAAGAAGCGATGGCGCGTCTGGGAAAACTCTTAGCTGCGG
AAGGCGGGCGTTGGACGGCCCTGTTAGAAGTGGTTGATCCGGAAGCGCCGGTCATGGTACCG
TATCAGGAACCAGGCGTGTATCTGCTGGCCCTCCGTTCGATTGGTGAAGGGCACTATCTTCT
GCCTGGGGTACATTTCCCGCTGCCTGAAGCCCTGCGTTACGTTCGGTGGGAACCACGCATGG
ACTTTGACCCTCATCGCTTTCGCGGTGAAATTCGCGACCTCCAAGGCGTAGAGGGCTACGTG
GTTACCGATGGTGCGGAGTTTGTCAAGTTCAAAACCGGCTGGGCGTTTCGGTTAGCGCGCTT
CCTGATGGACCCCGAAGGGGTGTTCCTGGAAGCCTATGCGGAAGATCGGCTGGACGACCTGG
TGGGTGCCTTGGCGGGCCGCGAGGACCTCCTGCGTGCGGTTGCGCGTGCGCAGGATTACCTG
GCAGGACTCTATGGTGAAGCAGTTGGAGCTGGCGATGCCTTACGCCGTATGGGCCTTCCGCG
CAAGGAAGCATGGGCGCGTGTACAGGAAGAGGCCGGTCGTTGGGGCGGCTTTGCCCCTGCGT
ATGCTCGCGCAGCAATGGCCGCATATGAAGGCGGCGAAGCCCGCGAAGCGTTTCTGGTCGAA
CTGCGCAAACGCTCCGCTCGTAAAGCTCTGGAAGCTCTGCACTTGTTCCCACGCGTTGGTGG
GGAATTACGCGGTGAATTC**TGGAGCCATCCGCAGTTTGAAAAA**

**Supplementary Information S2. Nucleotide sequence of a gene encoding codon-optimized human aprataxin.**

Recognition sequences for BamHI and EcoRI are underlined.

<u>GGATCC</u>ATGAGCAATGTGAATCTCAGCGTAAGCGATTTCTGGCGCGTGATGATGCGTGTCTG
TTGGTTGGTGCGGCAGGATTCACGCCATCAACGCATACGTCTGCCTCATTTGGAGGCTGTAG
TGATTGGCAGAGGTCCGGAAACGAAGATTACCGACAAAAAATGCAGTCGTCAGCAAGTCCAA
CTGAAAGCGGAATGCAACAAAGGGTACGTTAAGGTGAAACAGGTAGGTGTCAACCCCACGTC
CATTGATTCGGTGGTGATTGGGAAAGATCAGGAGGTAAAACTCCAACCAGGTCAGGTTCTGC
ACATGGTGAATGAGCTGTATCCGTATATCGTCGAGTTCGAAGAAGAGGCCAAAAATCCGGGC
CTCGAAACCCATCGCAAACGCAAACGTAGTGGCAACTCCGACTCTATCGAACGCGATGCAGC
CCAAGAAGCGGAAGCAGGTACTGGTCTTGAACCAGGCTCAAATAGTGGTCAGTGTTCGGTTC
CGCTCAAGAAAGGCAAGGATGCGCCGATCAAGAAAGAGAGTCTTGGCCATTGGTCCCAAGGC
TTGAAAATCTCCATGCAAGATCCGAAGATGCAGGTCTATAAAGACGAACAGGTTGTCGTGAT
CAAAGACAAATACCCGAAAGCGCGTTATCATTGGCTGGTTTTGCCCTGGACCAGCATTAGCT
CTCTGAAAGCCGTGGCACGTGAACACCTGGAGTTACTGAAACACATGCACACTGTAGGGGAA
AAAGTGATCGTTGACTTTGCGGGATCTTCGAAACTGCGCTTTCGCCTAGGCTATCACGCCAT
TCCGAGCATGTCACACGTGCATCTGCACGTTATTAGCCAGGATTTTGATTCACCGTGTCTGA
AGAACAAAAAGCATTGGAACAGCTTTAACACGGAATACTTCCTGGAATCTCAAGCTGTCATC
GAAATGGTGCAGGAAGCTGGTCGTGTTACCGTTCGAGACGGAATGCCTGAGCTGCTTAAATT
ACCGTTACGGTGCCATGAATGCCAGCAGTTACTGCCTTCGATTCCACAGCTGAAAGAACATC
TGCGTAAACATTGGACACAG<u>GAATTC</u>

**Supplementary Information S3.**

**Flowcharts for bioinformatic analyses**

### Trimming and mapping

**FASTQ**

↓ **fastp**: Remove low-quality reads

$ fastp (no option)

**FASTQ, filtered**

↓ **seqkit subseq** : Trim nucleobases added in TACS-T4

```
$ seqkit subseq -r 3:-2        #Read 1
                -r 2:-3        #Read 2
```

**FASTQ, trimmed**

↓ **bowtie2**: Map reads in paired-end mode

$ bowtie2 -q -p 10 -x *human_hg19*

**BAM**

↓ **samtools view**: Extract the reads whether mapped to nuclear genome, mapped to mitochondria, or unmapped

```
$ samtools view -f 2 | awk '$3!="LAMBDA"' |awk '$3!="chrM"’        #nuclear genome
$ samtools view chrM                                                #mitochondria
$ samtools view -f 4                                                #unmapped
```

**BAM**

↓ **awk**: Extract uniquely mapped reads

```
$ awk '{if(NR%2)ORS="~";else ORS="¥n";print}’ | grep -v ".*XS.*XS" |/
$ awk -F "~" '{ print $1 "¥n" $2 }’
```

**BAM, uniquely mapped**

### Separation of mapped reads and peak calling

#### BAM, uniquely mapped

**awk**: Separate the alignments by the fragment sizes

```
 $ awk '$9!=0’ | awk '{print sqrt($9^2) , $0}’ |/
  '$1 >= 35’ | awk '$1 <= 75’ | cut -f 2- -d " "       #C3D reads
  '$1 >= 147’ | awk '$1 <= 190’ | cut -f 2- -d " "   #NPD reads
```

#### BAM, extracted by fragment size

**samtools view**: Separate the alignments by the mapped reference genome strand

```
 $ samtools view “-f 128 -F 16” or “-f 80”          #reads mapped on top strand
 $ samtools view “-f 64 -F 16” or “-f 144”          #reads mapped on bottom strand
```

#### BAM, separated with mapped strand

**MACS2**: Peak call

$ macs2 callpeak -f BAMPE -g hs -q 0.01

$ cat *peak_top.bed peak_bottom.bed* > *peak_merged.bed*

#### BED

### Calculation of the read coverages

BAM

↓

**bedtools genomecov**: Calculate coverage of each fragments
$ bedtools genomecov -bga -pc

BEDGRAPH

↓

**bedGraphToBigWig**: Convert bedGraph to bigWig
$ bedGraphToBigWig (no option)

BIGWIG

### Size distribution plot
(Figures 2, 3, Supplementary Figures S5F-H, S6A)

**BAM**

**Calculation of size distribution from TLEN column in SAM format:**

```
$samtools view |cut -f 9 |tr -d "-” |/
$awk '{ v[$0]++ } END { for ( k in v ) print k "¥t" v[k]/2 }’ |/
$sort -n |awk '$1 <= 500'
```

**TSV**

Scatter plot made in Excel

**PDF**

### BLAST for unmapped reads
(Figure 3)

**BAM, unmapped to nuclear or mitochondrial genome**

↓ **SamtoFastq (picard tools):** Make fastq from bam
$ java -jar picard.jar SamToFastq (no option)

**FASTQ**

↓ **BBMerge (BBTools):** Merge paired fastq to single-end like fastq
$ bbmerge-auto.sh (no option)

**FASTQ, merged**

↓ **fastp:** Quality filtering of merged reads
$ fastp (no option)

**FASTQ, filtered**

↓ **fastq_to_fasta (fastx_toolkit) :** Convert fastq to fasta format
$ fastq_to_fasta (no option)

**FASTA**

↓ **BLASTn:** Realign of unmapped reads to microbial sequences
$ blastn -db *blastdb** -outfmt "17 SQ" /
-reward 1 -penalty -3 -word_size 12 -gapopen 5 -gapextend 2 -evalue 1E-4 /
-perc_identity 90 -culling_limit 2

* *blastdb* ;of blastdb_bacteria, blastdb_virus, blastdb_fungi
Source of the database sequences were downloaded from NCBI database (ftp://ftp.ncbi.nih.gov/genomes).

**BAM**

### Visualization with genome browser
(Figures 4A, 6A and 7C)

**BIGWIG,**
**BED**

↓

**UCSC genome browser:** Visualize the coverage and the peak tracks
Track Hub URL: http://itolab.med.kyushu-u.ac.jp/hisano/tracks_for_ucsc/hub.txt

**PDF**

### Comparing the coverage of C3D peaks among the donors (Figure 4C)

**BIGWIG**

↓

**multiBigwigSummary (deeptools)**: Calculate average coverage in equally sized bins
multiBigwigSummary bins --binSize=100

**NPZ, matrix file**

↓

**plotCorrelation (deeptools)**: Visualize the coverage correlation in bins
plotCorrelation --corMethod pearson --skipZeros --removeOutliers ¥
--whatToPlot scatterplot --xRange 0 500 --yRange 0 500

**PDF**

### Aggregation plot and heatmap
#### (Figures 4-7,Supplementary Figures S7-9, S11, S13-15)

**BED, regions**

**BIGWIG**

**Computematrix (deeptools):** Make matrix files for plot

```
$ computeMatrix reference-point --referencePoint center /
  --upstream 1000 --downstream 1000         #plot around the peak center
$ computeMatrix scale-regions ¥
  --upstream 1000 --downstream 1000         #regard the region range
```

**TXT, matrix file**

**plotHeatmap (deeptools):** Plot heatmaps

```
$ plotHeatmap
  --missingDataColor 'white' /
  --outFileSortedRegions
```

**PDF**

### HOMER peak annotation
#### (Figures 5A, 5B, Supplementary Figure S7D, S16)

**BED**

↓

**annotatePeaks (HOMER)**: Annotate peaks to known functional elements
$ annotatePeaks hg19 -annStats

**TSV**

↓

Bar chart made in Excel

**PDF**

### GO enrichment analysis
#### (Supplementary Figure S7B)

**TSV, result of HOMER annotation**

↓ Gene ID list of the C3D peaks with annotation "5' UTR " or "promoter" (n = 7,634) was selected according to the HOMER annotation.

**TXT, Gene ID list**

↓ **Panther gene list analysis** : Functional classification of genes
The analysis performed on website (http://pantherdb.org) and the result was downloaded as text file.
Total reference gene list supplied from the Panther website (n = 20,851).

**TXT**

↓ Fold enrichment was calculated against total reference genes in Excel

**R**: Fisher exact test of GO in C3D-regulated genes against reference genes.
fisher.test() (no option)

**PDF**

### Venn diagram, overlap test of the peaks
#### (Figures 6, 7, Supplementary Figures S6-10, S14)

**BED, regions**

**BED**

**ChIPpeakAnno:** Calculate and make venn diagram of overlapping peaks
findOverlapsOfPeaks() (no option)
makeVennDiagram() (no option)

**permTest (regioneR):** Permutation test of overlapping peaks

```
permTest(ntimes=1000, count.once=TRUE, genome="hg19",
         force.parallel=FALSE, randomize.function=randomizeRegions,
         evaluate.function=numOverlaps, verbose=FALSE)
```

**PDF**

### Calculation of the base composition of the peaks
#### (Figure 7B, Supplementary Figure S12)

**BAM**

↓

**bedtools bamtobed:** Acquire genomic coordinates from paired alignments
$ bedtools bamtobed -bedpe |awk 'BEGIN{OFS="¥t"} {print($1, $2, $6)}'

**BED**

↓

**bedtools getfasta:** Extract sequences from genome coordinates
$ bedtools getfasta (no option)

**FASTA**

↓

**seqkit fx2tab: C**alculate base contents of each fragments
$ seqkit fx2tab -g -G -B A -B T -B C -B G

**TSV**

↓

**ggplot2:** visualize histogram of base contents
$ ggplot(aes(x = **base*)) + geom_histogram(binwidth = 1)
**base*; A, T, C, G, GC-skew

Calculation of average base content in Excel

**PDF**

### Calculation of the peak strand bias of the peaks
#### (Supplementary Figure S12A)

**BED,**
**C3D**

**BIGWIG,**
**C3D separated by mapped strand**

**Computematrix (deeptools):** Make matrix files of coverage
$ computeMatrix scale-regions

**TXT, matrix file**

Strand bias of each C3D peak was calculated in Excel
strand_bias = (*Top* - *Bottom*) / (*Top* + *Bottom*)
*Top*; average coverages of the reads mapped on the top strand within C3D peak
*Bottom*; average coverages of the reads mapped on the bottom strand within C3D peak

**TSV**

**ggplot2:** Visualize histogram of base contents
$ ggplot(aes(x = strand_bias)) + geom_histogram(binwidth = 1)

**PDF**

### G4motif calculated by the Quadparser algorithm
#### (Figure 7 and Supplementary Figures S14-16)

**FASTA, whole genome**

**fastaregexfinder**: Search a fasta file for matches to a regex and return a bed file
(https://github.com/dariober/bioinformatics-cafe/blob/master/fastaRegexFinder/)
fastaRegexFinder.py -r '([gG]{3,}¥w{1,7}){3,}[gG]{3,}'

**BED**

### Random coordinates from hg19 genome
#### (Figure 7, Supplementary Figures S9, S10, S12)

**FASTA, whole genome**

**bedtools random**: Generate a random set of intervals from reference genome
$ bedtools random -seed 11 -l 55 -n 100000

**BED**
